# Characterizing structural and kinetic ensembles of intrinsically disordered proteins using writhe

**DOI:** 10.1101/2025.04.26.650781

**Authors:** Thomas R. Sisk, Simon Olsson, Paul Robustelli

## Abstract

The biological functions of intrinsically disordered proteins (IDPs) are governed by the conformational states they adopt in solution and the kinetics of transitions between these states. We apply writhe, a knot-theoretic measure that quantifies the crossings of curves in three-dimensional space, to analyze the conformational ensembles and dynamics of IDPs. We develop multiscale descriptors of protein backbones from writhe to identify slow motions of IDPs and demonstrate that these descriptors provide a superior basis for constructing Markov state models of IDP conformational dynamics compared to traditional distance-based descriptors. Additionally, we leverage the symmetry properties of writhe to design an equivariant neural network architecture to sample conformational ensembles of IDPs with a denoising diffusion probabilistic model. The writhe-based frameworks presented here provide a powerful and versatile approach for understanding how the structural ensembles and conformational dynamics of IDPs influence their biological functions.

**Significance Statement:** Intrinsically disordered proteins (IDPs) are essential for many cellular processes and are implicated in numerous diseases. The biological functions of IDPs are dictated by the populations of the diverse conformational states they adopt in solution and the kinetics of the conformational transitions between these states. Computer simulations are a powerful tool for studying IDPs, but analyzing simulated structural ensembles of IDPs to identify functionally important conformational states and motions remains a significant challenge. In this study, we demonstrate that writhe, a geometric descriptor from the field of knot theory that describes the crossing of curves in space, provides a powerful basis for characterizing the structural ensembles and conformational dynamics of IDPs in atomic detail. Our results provide new tools for unraveling the relationships between IDP sequences, their conformational ensembles, and their biological functions.

## Introduction

Intrinsically disordered proteins (IDPs) populate heterogeneous conformational ensembles of interconverting structures in solution and comprise approximately one-third of the human proteome.^1^ While the physiological interactions and cellular functions of folded proteins are largely determined by their three-dimensional (3D) structures, the biological functions of IDPs are dictated by the properties of the dynamic conformational ensembles they adopt in solution and when bound to their physiological interaction partners.^2–8^ The physiological interactions of IDPs are determined by the populations of the conformational states they adopt in solution (their *structural ensembles*), the kinetics of the conformational transitions between these states (their *kinetic ensembles*), and the thermodynamics and kinetics of their binding events and folding-upon-binding pathways. There has been substantial progress in efforts to characterize the structural ensembles of IDP at atomic resolution.^2–5^ Methods to determine atomic resolution kinetic ensembles of IDPs, which describe the structures, populations, and interconversion rates of IDP conformational states, have only recently begun to emerge.^6, 7^

Due to their highly dynamic nature, characterizing structural and kinetic ensembles of IDPs in atomic detail with biophysical experiments is extremely challenging and generally requires integrating biophysical experiments with all-atom molecular dynamics (MD) computer simulations.^4, 7, 8^ Advances in the accuracy of physical models, or *force fields*, used in all-atom MD simulations have dramatically enhanced the reliability of atomistic IDP ensembles.^2, 3, 5, 9, 10^ Identifying kinetically distinct conformational states of IDPs, however, remains a substantial challenge. Markov state models (MSMs), which describe the dynamics of stochastic systems as a transition network of memoryless, probabilistic jumps between conformational states, are a promising approach for building kinetic ensembles of IDPs from MD simulations.^11–14^

Building accurate MSMs of IDPs requires identifying molecular features that describe the slowest structural fluctuations observed in MD simulations and using these features to partition MD trajectories into discrete, metastable states. As IDPs have a large number of degrees of freedom, their conformational space is extremely high-dimensional, and identifying slowly evolving structural features to partition IDP trajectories into structurally and kinetically distinct conformations is challenging.^6, 7^ The variational approach to Markov processes (VAMP) provides a powerful theoretical framework to identify slowly evolving molecular features in MD simulations quantitatively.^15–20^ The VAMP method, which is based on time-lagged canonical correlation analysis (tCCA)^15, 21, 22^, uses a family of dimensionality reduction methods and variational scores to identify slowly varying collective variables among a collection of candidate features and transform these features into slowly evolving, low-dimensional reaction coordinates. VAMP methods have proven highly valuable for building MSMs from biomolecular simulations.^15–18, 20^

General and robust sets of molecular features that effectively describe the conformational dynamics of IDPs have yet to be identified. Due to the heterogeneity of IDP conformational spaces and their highly diffusive dynamics, many conventional molecular features used to characterize kinetic ensembles and build MSMs of structured proteins are ineffective for IDPs. Fluctuations of similarity measures to 3D reference structures (such as RMSD), dihedral angles, Euclidean interatomic distances, and secondary structure order parameters often fail to meaningfully separate IDPs into kinetically distinct conformational states, as these properties can fluctuate within conformational substates of IDPs on fast nanosecond timescales. Global order parameters that fluctuate on longer timescales, such as the radius of gyration or total solvent accessible surface area of IDP conformations, are often too coarse to identify conformational states of IDPs at the fine-grained resolution required to provide insight into their physiological interactions and biological functions.

The fields of knot theory^23, 24^ and differential geometry^25, 26^ offer promising alternatives to traditional molecular features for identifying discrete, metastable conformational states of IDPs and characterizing their transition kinetics. The geometric descriptor *writhe*, which quantifies the orientations of crossings of curves in 3D space, has previously been applied to compare the conformations of folded proteins^27–31^ and characterize the coiling of DNA.^28^ Here, we demonstrate that the writhe of protein backbones provides a powerful basis for characterizing the structural ensembles and conformational dynamics of IDPs.

In this study, we develop descriptions of the writhe of protein backbones on multiple length scales. We show that these descriptors capture distinct structural properties with unique relaxation timescales and form a general and robust basis for constructing atomic-resolution kinetic models of IDP conformational dynamics. We use multiscale writhe descriptors to build MSMs from long timescale all-atom MD simulations of several IDPs and a fast-folding protein, and compare these to MSMs derived using traditional Euclidean distance features. We find that writhe descriptors identify more kinetically and structurally distinct conformational states than traditional distance features and that MSMs built from writhe descriptors capture more kinetic variance and resolve longer timescale processes than MSMs built from distance descriptors for all systems examined in this study. We use multiscale writhe descriptors to build an MSM of the conformational dynamics of the intrinsically disordered Aβ42 peptide from a large collection of previously reported MD simulations.^7^ Our analysis demonstrates that the kinetic metastability of Aβ42 conformational states can be intuitively understood in terms of the relative orientations of backbone chain crossings. Together, these results demonstrate that the writhe descriptors presented here provide a powerful basis for describing the conformational dynamics of IDPs observed in molecular simulations.

Generative artificial intelligence (AI) is an emerging alternative approach to modeling conformational ensembles of proteins at substantially reduced computational cost.^32–35^ Instead of explicitly simulating physical motions, as in MD simulations, generative AI models learn from data (e.g., experimental structures, protein sequences, or MD trajectories) to predict unknown structures directly from protein sequences. Recent breakthroughs in AI-driven protein structure prediction, such as AlphaFold, are revolutionizing the computational modeling of folded proteins and other systems characterized by single structures.^36, 37^ Notable works aimed at sampling ensembles of structures include Boltzmann generators^35^, which utilize molecular dynamics force fields to generate structures and their Boltzmann weights, and implicit transfer operators, which learn to advance the state of a system over variable time steps to overcome long timescale barriers that hinder sampling in simulation.^34^ Recent applications of deep generative techniques to IDPs show substantial promise^32, 38, 39^ , but many methodological questions remain open.

It is currently unclear what generative AI model architectures, input features, and training strategies will most efficiently produce physically realistic IDP ensembles. Recent studies^34, 40, 41^ have shown that neural networks trained to sample protein conformations in generative models can be made substantially more robust by satisfying relevant symmetry constraints. For MD simulation data, it is highly desirable for such neural networks to have the property of SE(3)-equivariance, meaning the neural network responds predictably when input structures are rotated or translated. SE(3)-equivariant neural network architectures ensure that distributions of conformations from generative models are not affected by global rotations and translations of molecular structures.^40^ Another important property of SE(3)-equivariant all-atom generative models of protein structures is that they do not invert the chirality of L-amino acids and D-amino acids in generated structures.^34^ Here, we show that the orientations of IDP chain crossings in one-particle-per-residue representations of IDPs popular in coarse-grained simulations and generative models^39, 42, 43^ also exhibit chirality. We demonstrate that IDP chain crossings with mirror image reflected orientations have oppositely signed writhe (i.e., writhe is a *parity-odd pseudoscalar*). We leverage this symmetry property of writhe to design an efficient SE(3)-equivariant neural network to sample IDP conformations with a score-based denoising diffusion probabilistic model^44^ (DDPM) and present a proof-of-principle demonstrating this architecture can be used to accurately reproduce IDP conformational distributions obtained from MD simulations.

## Results

### Calculating the writhe of protein conformations

The field of knot theory studies the geometry, deformation, and equivalence of closed curves in three dimensions (3D).^45, 46^ The central challenge in knot theory is to determine whether two knots are equivalent, or *isotopic*. Equivalence is confirmed by finding a set of deformations that map one knot to another without breaking or passing through itself.^45, 46^ Many ideas and mathematical descriptions from knot theory can be used to characterize conformational states of polymers, given that many of their conformational transitions are governed by similar principles.^47, 48^ Mathematical knots are commonly represented via *knot* diagrams, where a 3D curve is projected onto a 2D plane and drawn to preserve oriented crossings. By specifying the directionality of the curve, one can designate oriented crossings as positive or negative. The total *writhe* of a knot diagram can be computed as the sum of its *signed* (or oriented) crossings. For a continuous curve in 3D, the writhe can be expressed as the Gaussian integral^23, 28^:

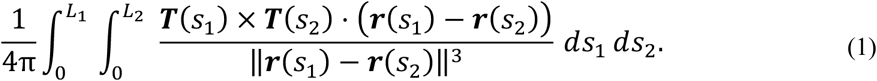

Here, 𝒓(𝑠) represents the position vector of a point along the curve parameterized by the arc length, 𝑠, which takes values in the interval [0, L]. This interval represents the entire length of the curve, L. The function 𝑻(𝑠) is the unit tangent vector at 𝑠, defined as 𝑻(𝑠) =𝑑𝑟⁄𝑑𝑠, which describes the local direction of the curve. The parameters 𝑠_1_ and 𝑠_2_ serve as integration variables, allowing the curve to be integrated over itself to account for all possible pairs of points that contribute to the writhe. Additional discussion of the calculation of Gaussian integrals of continuous curves is included in the Supplementary Information section “Gaussian integrals and writhe of continuous curves”.

To compute the writhe of a protein conformation, an open polygonal curve can be constructed from normalized displacement vectors between atoms along the backbone, resulting in a set of *segments* (Figure 1). These segments serve as finite approximations of the tangent vectors 𝑻(𝑠) in Equation 1. One possible segmentation is to describe the protein backbone as a series of segments connecting consecutive Cα atoms (i.e., vectors from Cαᵢ to Cαᵢ₊₁).^28, 29^ After segmenting the curve into a finite number of elements, the writhe can be computed pairwise between all segments, and the resulting set of crossings can be organized into a symmetric matrix that we refer to as the *writhe matrix* (Figure 1D). In the discrete formulation, the writhe is determined from the relative orientations of segments, which implicitly depend on their spatial separations (Equation 1, Figure 1, Supplementary Figure S1). As a result, the writhe matrix resembles a contact map but also encodes the relative orientation between each pair of segments. Further discussion of the numerical computation of the writhe from line segments is provided in Supplementary Information Appendix A, “Numerical computation of the writhe and algorithms.”

**Figure 1.**
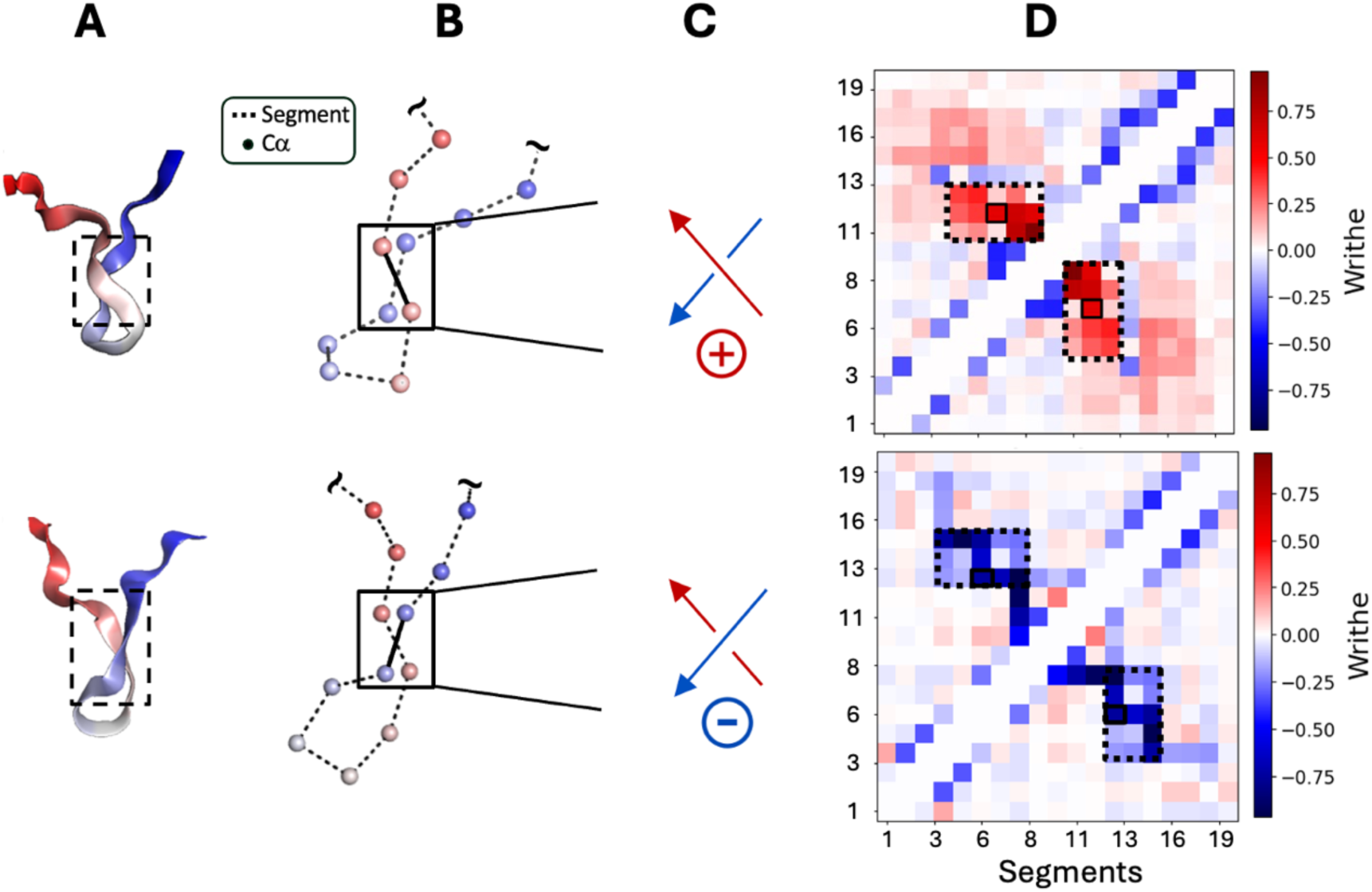
Computing the writhe of protein conformations. **(A)** Two conformations sampled from an unbiased, long-timescale equilibrium MD simulation of a 20-residue fragment of α-synuclein^69^ that exhibit backbone chain crossings with opposite-signed writhe. Structural representations of the α-synuclein fragment are colored with a blue-to-red gradient from the N-terminus to the C-terminus. **(B)** Illustration of backbone segments constructed from displacement vectors between neighboring (Cα_i_-Cα_i+1_) Cα atoms. **(C)** The sign and handedness of the segment crossings enclosed with solid black boxes in panel B. **(D)** A symmetric “*writhe matrix”* displaying the pairwise writhe values between all Cα_i_-Cα_i+1_ segments for the conformations displayed in (B). The matrix indices enclosed by dashed lines correspond to the writhe of segments contained in the region marked with a dashed box in panel (A).

Here, we use a geometric approach to compute the writhe of a chain defined by discrete segments.^28, 29^ We evaluate the integral in Equation 1 for individual pairs of segments by computing a solid angle that quantifies their apparent crossing from all viewpoints in space.^28^ We visualize the computation of the writhe with this geometric approach for a single pair of segments in Figure 2. Figure 2B illustrates that this computation is equivalent to computing the surface area of a spherical quadrilateral enclosed by vertices defined by the relative orientation of the crossing segments as seen from the perspective of each view direction vector, 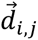. In Appendix A in the Supplementary Information, we provide an overview of existing algorithms for the numerical calculation of writhe and introduce a new algorithm to efficiently compute the writhe with reduced wall-clock times (Supplementary Table 1).

**Figure 2.**
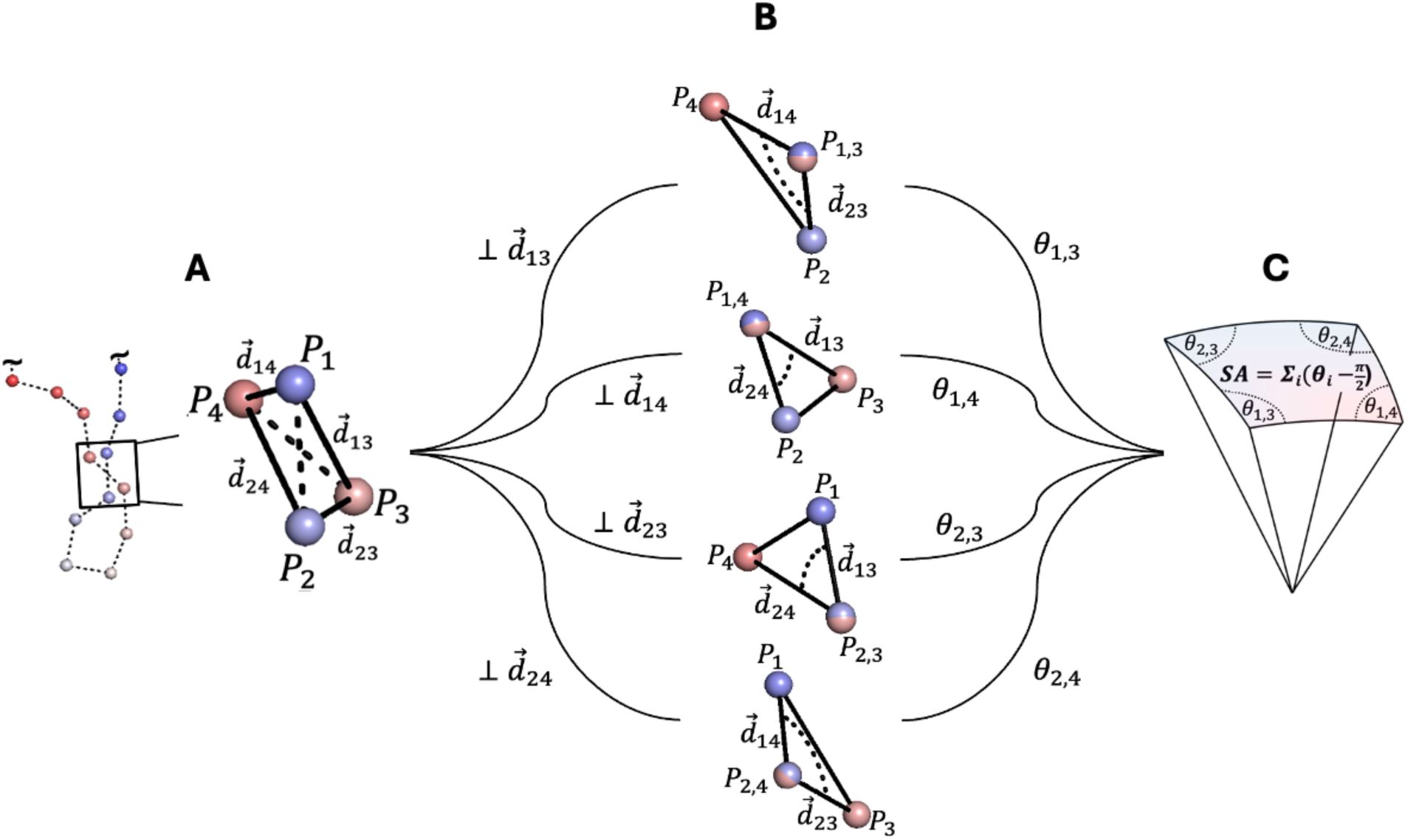
Geometric computation of writhe between two backbone segments of a protein. **(A)** Cα atoms of a protein backbone conformation are shown as spheres and colored with a gradient from the N-terminus (blue) to the C-terminus (red). Cα_i_-Cα_i+1_ segments defining the protein’s backbone trace are shown as dashed lines. The segment crossing enclosed in a black box is magnified to the right, showing the view direction vectors 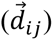 between the endpoints of the segments used in the computation of the writhe (shown as solid black lines). The writhe of a pair of discrete segments is defined as the summation of apparent crossings as observed from the perspective of each of the four view direction vectors, 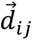. **(B)** Projecting orthogonally to each 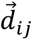 generates a view where the points 𝑃_𝑖_ and 𝑃_*j*_ appear to coincide (𝑃_𝑖,*j*_, red-blue points), and two corresponding view direction vectors create a vertex that admits the angle, 𝜃_𝑖,*j*_. The four vertices defining a spherical quadrilateral (shown in C) are shown from the perspective of each view direction, 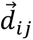 **(C)** Placing each view direction vector at the origin and its associated vertex on the surface of the unit sphere constructs a spherical quadrilateral (or quadrangle). The surface area (SA) of the quadrangle, normalized by 2𝜋 is equal in magnitude to the writhe.

### Characterizing protein conformations using writhe at multiple length scales

The writhe of a protein backbone can be computed at multiple length scales. In previous studies, segments have predominantly been constructed from displacement vectors between adjacent Cα atoms (Cα_i_-Cα_i+1_) (Figure 1).^29–31, 47, 48^ A previous approach to obtain higher order writhe descriptors of protein structures was introduced by Rogan et al., who investigated higher order Gaussian integrals inspired by Vassiliev knot invariants to identify similarities between the global fold structures of proteins to classify them.^30, 47, 49^ We develop multiscale writhe descriptors by simultaneously analyzing the writhe of protein conformations using multiple *segment lengths*. Here, the segment length *l* specifies the offset of Cα atoms (Cα*_i_*-Cα*_i_*_+*l*_) used to define segments in a writhe calculation. Increasing the segment length effectively smooths the polygonal curve representing the protein’s backbone.^31^ This reduces signal from local backbone crossings, such as the presence of secondary structure, and more effectively captures longer length scale structural features and fluctuations (Figure 3).

**Figure 3.**
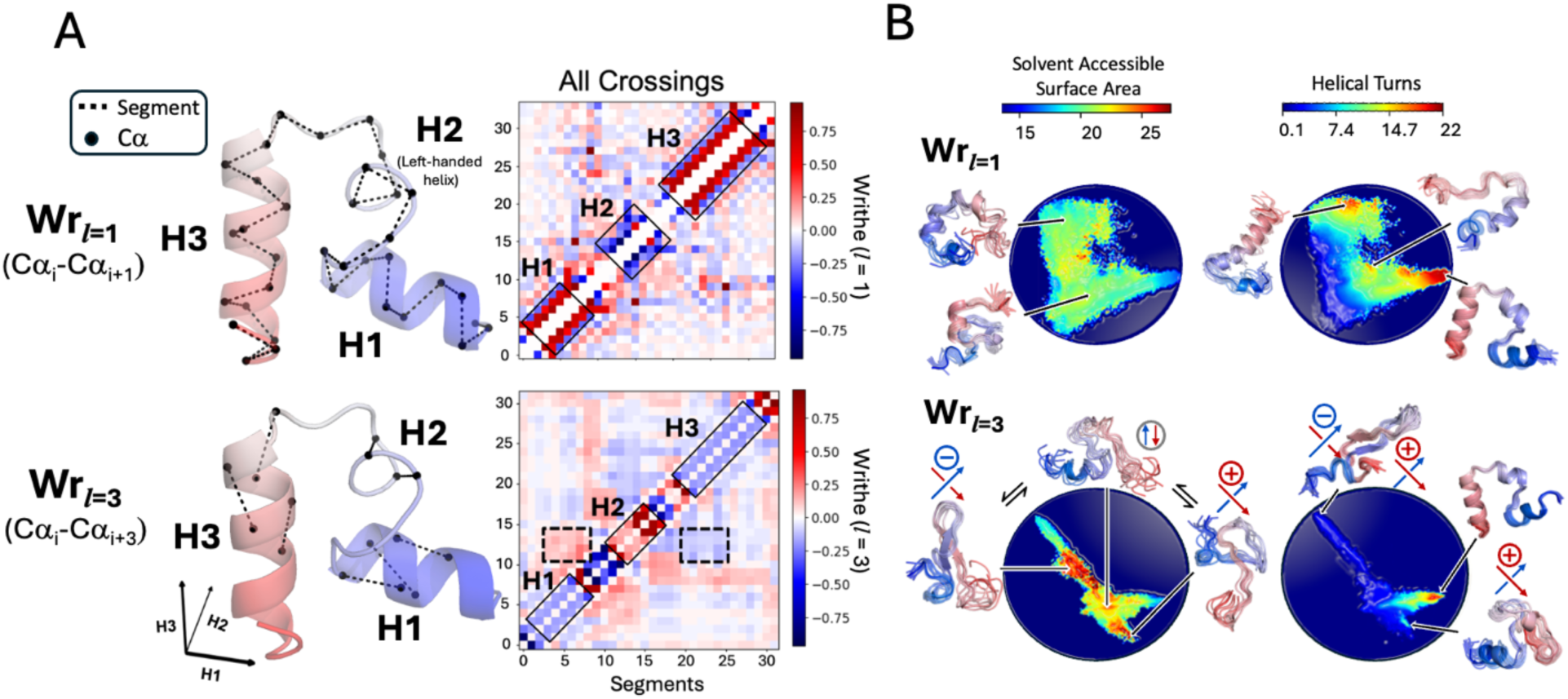
Describing geometric properties of proteins at different length scales with writhe. **(A)** The writhe of a conformation taken from an unbiased, long-timescale (319 μs) equilibrium MD simulation of wild-type HP35 calculated from segments between adjacent Cα atoms (Wr*_l_*_=1_) and every third Cα atom (Wr*_l_*_=3_). Structures are colored with a blue (N-terminus) to red (C-terminus) gradient and are shown with select segments between Cα atoms used in the computation of the writhe as black, dashed lines. The matrices of all pairwise contributions to the writhe are shown to the right of each structure, with segments corresponding to the H1, H2, and H3 domains highlighted along the diagonal with solid black ones. In the Wr*_l_*_=3_ writhe matrix, off-diagonal elements reflecting the relative orientations of the H2 domain with H1 and H3 domains are highlighted with dashed lines. **(B)** Projections of each simulation frame onto the two dominant time-lagged canonical components obtained from performing time-lagged canonical correlation analysis (tCCA) on writhe descriptors computed from Wr*_l_*_=1_ and Wr*_l_*_=3._ Projections are colored by the solvent-accessible surface area and the alpha helical order parameter, Sα^51^. Representative structures are shown adjacent to the projections with the handedness of the crossing demarcated where relevant.

To denote the segment length (*l*) used to compute a set of writhe features, we adopt the shorthand notation Wr*_l_*. Wr*_l=_*_1_ features correspond to writhe features computed from (Cα_i_-Cα_i+1_) segments, while Wr*_l=_*_3_ features correspond to writhe features computed from (Cα_i_-Cα_i+3_). We illustrate the geometric differences in writhe features computed from segment lengths *l*=1 (Wr*_l_*_=1_) and *l*=3 (Wr*_l_*_=3_) for conformations of the fast-folding protein, HP35, in Figure 3. Figure 3A shows a representative conformation of HP35, obtained from a previously published 319 μs MD simulation^50^, depicted with segments of length *l*=1 and *l*=3. The corresponding Wr*_l_*_=1_ and Wr*_l_*_=3_ writhe matrices for this conformation are also presented. This conformation contains three helical domains, H1, H2, and H3. H1 and H3 are right-handed helices, and H2 contains a left-handed helical turn. The handedness of the helices is resolved by the sign of the writhe features computed at Wr*_l_*_=1_ (Figure 3A). In contrast, the Wr*_l_*_=3_ matrix shows reduced fluctuations in the values of the writhe of neighboring segments, and more effectively captures the relative orientations of the helical domains, seen as off-diagonal elements in the Wr*_l_*_=3_ writhe matrix (Figure 3A).

In Figure 3B, we visualize the results of time-lagged canonical correlation analysis^15^ (tCCA; see Methods) applied to writhe features computed for all frames of the 319μs simulation of HP35. The analysis was performed using either Wr*_l_*_=1_ or Wr*_l_*_=3_ features; we compare projections of the HP35 MD trajectory onto the two slowest evolving time-lagged canonical components (tCCs) obtained with each segment length. We characterize the 2D tCCA projections using 2D histograms colored by the average values of the α-helical order parameter Sα^51, 52^ and the solvent accessible surface area of all the conformations in each bin. We observe that the tCCA projection of Wr*_l_*_=1_ writhe features is sensitive to the presence of local secondary structure in HP35, and clearly separates states based on the number and location of canonical helical elements, as quantified by the α-helical order parameter Sα.^51^ In contrast, the Wr*_l_*_=3_ tCCA predominantly captures more global chain rearrangements with larger differences in the distribution of solvent accessible surface area (SASA). Representative structures from high SASA regions in the Wr*_l=_*_3_ tCCA projection exhibit delocalized crossings with differing orientations involving residues distant from each other in sequence (Figure 3B). These results demonstrate that writhe features computed at different length scales are sensitive to distinct conformational rearrangements, motivating our use of multiscale writhe descriptors to build kinetic models of IDP conformational dynamics.

### Characterizing the conformational dynamics of intrinsically disordered proteins using multiscale writhe descriptors

To assess the ability of multiscale writhe descriptors to characterize IDP conformational states and elucidate slow dynamic modes, we compute the writhe using several segment lengths (*l*) for a diverse set of previously published long-timescale molecular dynamics simulations. This simulation dataset includes long timescale equilibrium MD simulations of four IDPs performed with the a99SB-*disp* protein force field and a99SB-*disp* water model^2^: a 73μs simulation of α-synuclein (140 residues)^2^, 30 μs simulations of the partially helical IDPs ACTR (71 residues)^2^ and PaaA2 (71 residues)^2^, and a 100 μs simulation of the α-helical molecular recognition element of N_TAIL_ (21 residues, which we subsequently refer to as “N_TAIL_”)^52^. We also analyze a 319μs simulation of wild-type HP35 (35 residues), the fast-folding Villin headpiece subdomain^50^, performed with the amber ff99SB*-ILDN^53, 54^ protein force field and TIP3P water model^55^ and a collection of 5120 independent MD simulations (with an aggregate simulation time of 315μs) of Aβ42 (42 residues) performed with the CHARMM22* protein force field^56^ and the TIP3P water model. These trajectories were selected based on their excellent agreement with experimental data, as reported in their original publications.^2, 7, 50, 52^

For each MD trajectory, we apply tCCA to writhe features computed at different segment lengths and apply tCCA to inter-residue distances and compare the resulting kinetic variances (Equation 3). The kinetic variance of tCCA is equivalent to the “VAMP-2 score”^15–17^, a metric used in the variational approach to Markov processes (VAMP) framework that quantifies how well a set of features captures the slowest timescale dynamics of a system.^20^ A larger kinetic variance indicates that a set of features is better suited for MSM construction. In Supplementary Figure 2, we compare the kinetic variance captured by the 10 largest tCCA components of writhe features computed with different segment lengths and inter-residue distances for each MD trajectory. We evaluate writhe features computed at single segment lengths (Wr*_l=_*_1_, Wr*_l=_*_3_, or Wr*_l=_*_5_). We also applied tCCA to concatenated writhe features computed at multiple segment lengths, which we refer to as “multiscale writhe descriptors”. The multiscale writhe descriptors considered here include Wr*_l=_*_1,3,_ Wr*_l=_*_1,2,3,_ and Wr*_l=_*_1,3,5_ (where Wr*_l=_*_1,3_ indicates that both Wr*_l=_*_1_ and Wr*_l=_*_3_ features were used as inputs for tCCA). For each system examined, the multiscale writhe features, Wr*_l=_*_1,2,3_ and Wr*_l=_*_1,3,5_, capture the greatest kinetic variance, indicating that these multiscale representations more effectively describe longer-timescale kinetic processes.

For additional insight, we compare the autocorrelation times of writhe features computed at different length scales for α-synuclein, ACTR, PaaA2, HP35, and N_TAIL_ in Supplementary Figure 3. We find that writhe features computed at longer segment lengths are less sensitive to structural fluctuations at short length scales and are more sensitive to structural fluctuations between segments more distant in sequence. In contrast, we observe that writhe features computed at segment length *l=*1 excel at capturing local structural features like α-helices (Figure 3). Taken together, our results show that multiscale writhe descriptors effectively describe long timescale structural fluctuations of IDPs that are not well described by Euclidean distances or writhe computed at a single length scale.

### Building Markov state models of intrinsically disordered proteins using writhe

We illustrate the impact of incorporating writhe in the construction of kinetic models by comparing MSMs built from writhe to MSMs built from inter-residue distances for a 30 μs simulation of ACTR (see Methods). To enable a direct comparison of writhe input features and distance input features with similar dimensions, we first performed tCCA and constructed MSMs from the ACTR MD trajectory using inter-residue distances and writhe features computed at a single segment length, Wr*_l=_*_1_. We compare the properties of the tCCA projections obtained from writhe and from distances in Figure 4. We project the free energy surface of the ACTR MD trajectory onto the two slowest evolving tCCA components obtained from Wr*_l=_*_1_ or inter-residue distances, respectively, in Supplementary Figure S4. We observe that the slowest evolving tCCs obtained from distances and Wr*_l=_*_1_ resolve similar numbers of states, however, there is a large difference in the number of distinct states resolved by the second slowest evolving tCCs (Figure 4A). We observe that the second slowest evolving tCC obtained from Wr*_l=_*_1_ isolates several additional free energy basins compared to the second largest tCC obtained from inter-residue distances. We quantify the structural resolution of each coordinate for a series of K-means^57^ cluster assignments using the silhouette score^58^ (a measure of the consistency of a clustering) (Figure 4B). We observe that the silhouette scores for the distance and writhe tCCA projections are maximized at *k* = 3 and *k* = 6 K-means clusters, respectively. We observe that the silhouette score of K-means clusters obtained from writhe doesn’t significantly decline until *k* = 10 clusters, while the silhouette score of K-means clusters obtained from inter-residues substantially declines after *k* = 3 clusters. These results demonstrate that fluctuations in Wr*_l=_*_1_ resolve substantially more distinct states than fluctuations of inter-residue distances.

**Figure 4.**
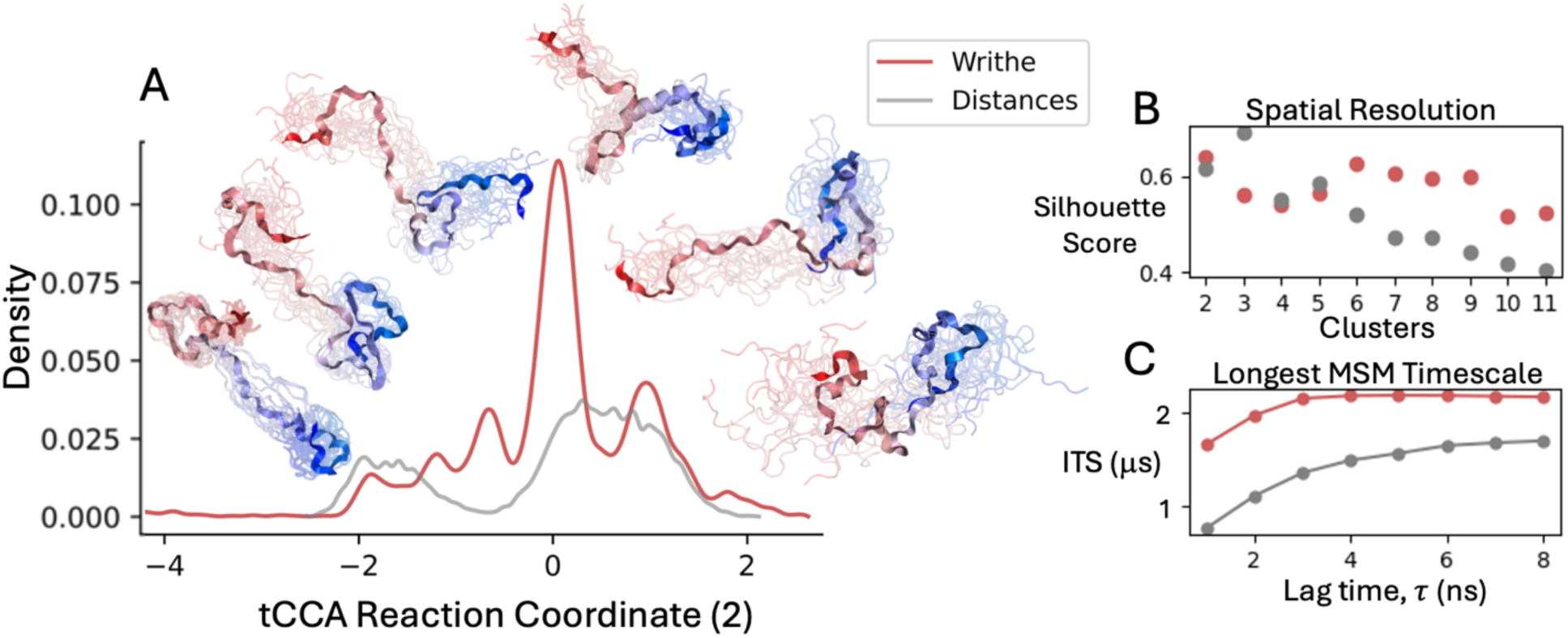
Comparing reaction coordinates, states, and MSM observables derived from writhe and Euclidean distances. **(A)** Reaction coordinates obtained from time-lagged canonical correlation analysis (tCCA) on writhe features computed using segments obtained from adjacent Cα atoms (Wr*_l_*_=1_) (red) and Euclidean distances between all Cα atoms (gray), for all frames from a continuous 30 μs equilibrium MD simulation of the intrinsically disordered protein ACTR. **(B)** Silhouette scores, reflecting the quality of cluster assignments, as a function of the number of K-means clusters applied to the one-dimensional reaction coordinates.^58^ **(C)** The longest implied timescales obtained from Markov state models (MSMs) constructed by clustering the first three dominant time-lagged canonical components from each dataset using 40 K-means clusters.

We proceed to estimate 40-state MSMs of ACTR by applying K-means clustering on the three slowest evolving tCCA components obtained from writhe and inter-residue distances, independently (see Methods). We compare the maximum likelihood^59^ estimates of the largest implied timescales (ITS) of these MSMs as a function of model lag time in Figure 4C. The largest ITS describes the timescale of the slowest processes captured by an MSM. We observe that the largest ITS of the Wr*_l=_*_1_ MSM converges to a substantially larger value (∼ 2.0 μs) than the largest ITS of the distance MSM (∼0.375 μs). The largest ITS of the writhe MSM also converges at shorter lag times. This demonstrates that the ACTR MSM constructed from Wr*_l=_*_1_ input features captures slower dynamic processes than an MSM constructed from inter-residue distances. We compute an additional ACTR writhe MSM using the Wr*_l=_*_1,3,5_ feature set, which was found to capture the most kinetic variance of the ACTR MD simulation (Supplementary Figure S2). We estimate an MSM using the 3 slowest evolving tCCs, 40 K-means clusters, and a lag time of 6 ns (see Methods). We coarse-grain the ACTR MSM to 7 macro-states using PCCA++ spectral clustering^60–62^ and display MSM validation metrics in Supplementary Figure S5.

We next use Wr*_l=_*_1,3,5_ multiscale writhe descriptors to build an MSM from a previously reported set of 5120 independent MD simulations (315 μs of cumulative simulation time) of the Alzheimer’s disease-associated peptide Aβ42.^7^ These simulations were previously used to construct an MSM using the deep learning VAMPnet approach with inter-residue distance inputs.^7, 63, 64^ We performed tCCA on this simulation dataset using Wr*_l=_*_1,3,5_ descriptors and inter-residue distances as inputs (see Methods). We compare projections of the Aβ42 trajectories onto the two slowest evolving tCCA components obtained from writhe and the two slowest evolving tCCA components obtained from distances in Supplementary Figure S6. We observe that the distance tCCA projections resolve four free-energy basins, while the Wr*_l=_*_1,3,5_ tCCA projection resolves several additional free-energy basins.

We proceed to construct an MSM from the extensive Aβ42 MD simulation dataset and find that we can construct a valid 5-state MSM using the multiscale Wr*_l=_*_1,3,5_ writhe descriptors (see Methods). We present a visual depiction of the conformational ensembles and the average writhe matrices of the five metastable conformational states in a MSM transition network in Figure 5. We present MSM validation metrics of the Aβ42 writhe MSM in Supplementary Figure S7. We display the macro-state transition flux matrix and mean first passage time (MFPT) matrix of the Aβ42 writhe MSM in Supplementary Figure S8 and Supplementary Figure S9, respectively. Figure 5 illustrates that the kinetic separation of the metastable states observed in simulations of Aβ42 can be intuitively understood by the orientation of long-range contacts in each state. Comparison of the equilibrium-weighted, average writhe matrices of states 4 and 2 illustrates that their long-range contacts have opposite crossing orientations. Consequently, there is little transition flux between these states directly, and interconversions primarily proceed through state 1, the most disordered and extended metastable state.

**Figure 5.**
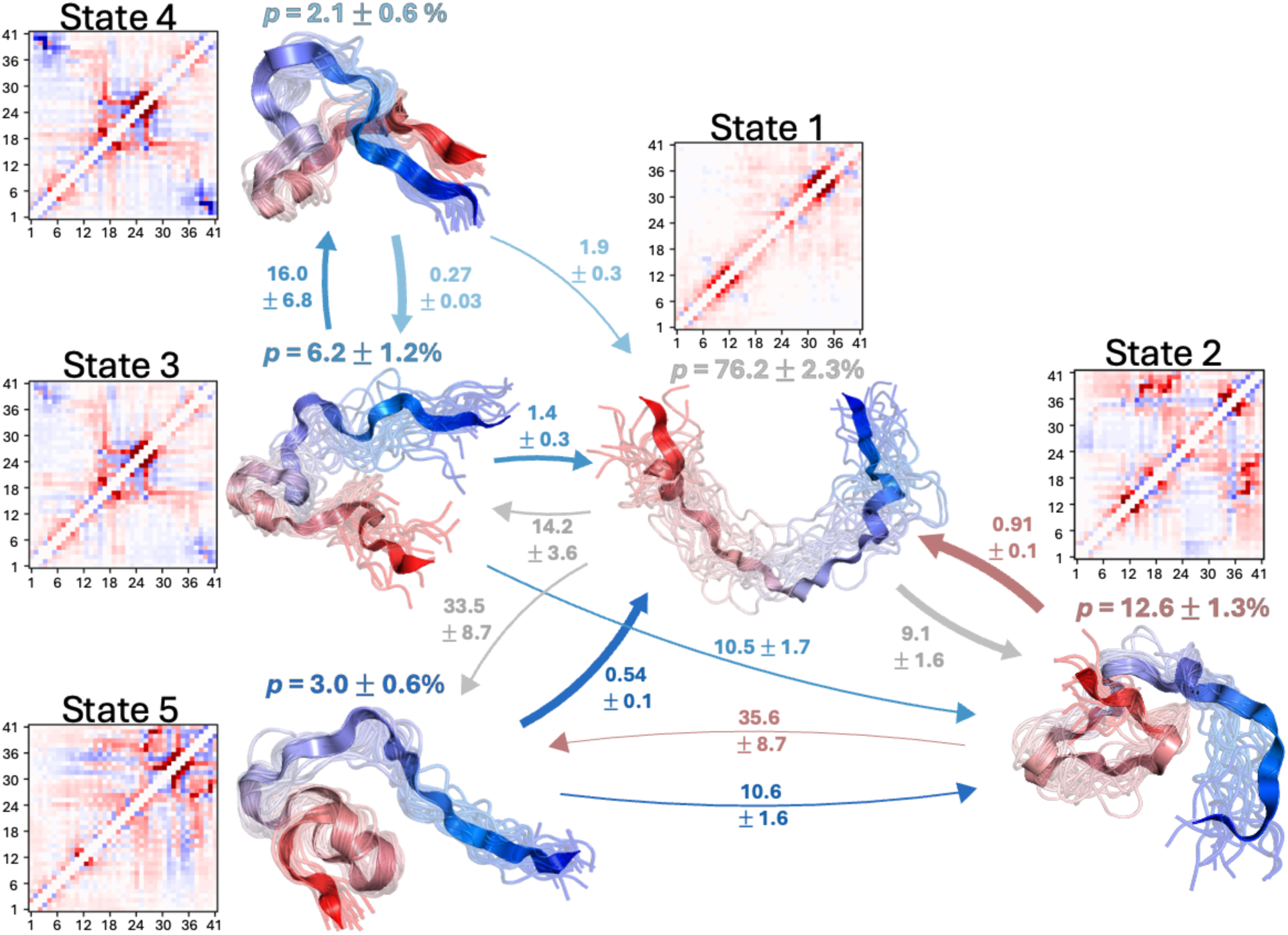
Markov state model (MSM) of Aβ42 derived constructed from multiscale writhe descriptors. Transition network representation of the transition probabilities and transition rates obtained from a coarse-grained MSM derived from 315 μs of MD simulations of Aβ42 using multiscale writhe features. Representative structures of each Markov state are displayed in circles along with their stationary probabilities (*p*). In representative structures of each state, Aβ42 is colored with a blue-to-red gradient from the N-terminus to the C-terminus. Transition probability fluxes between states are indicated with directed arrows, and the thickness of the arrows is proportional to the magnitude of the flux between states. Mean first passage times between states are reported in μs. All errors indicate the mean of the upper and lower deviations of the 95% confidence interval calculated from bootstrapping using 1000 samples.

To further demonstrate the ability of writhe descriptors to describe the conformational dynamics of IDPs, we systematically compare the properties of MSMs estimated from writhe and distances for α-synuclein, ACTR, PaaA2, Aβ42, HP35, and N_TAIL_ (see Methods). For MD simulations of each protein, we perform tCCA on inter-residue distances and perform tCCA on multiscale writhe features. For each system, we identify the set of writhe features that produce the largest kinetic variance obtained from tCCA (Supplementary Figure S2) and use this writhe feature set to construct MSMs. We compare MSMs constructed from the selected writhe feature set with MSMs constructed from inter-residue distances using several combinations of tCCA projections and numbers of clusters. We apply K-means clustering to 2, 3, 5, and 10-dimensional tCCA projections and estimate MSMS using 10, 20, and 40 cluster centers. In Supplementary Figure S10, we compare the convergence of the largest implied timescale of each MSM estimated with grid scans over the number of cluster centers and tCCA dimensions as a function of the MSM lag time. We observe that MSMs built using writhe produce longer implied timescales (describing slower processes) with substantially improved convergence for all simulation datasets.

### Using writhe to construct a generative model of IDP ensembles

There is a growing interest in developing generative models to predict the conformational ensembles of IDPs directly from sequence.^32, 36–38, 65^ Modeling protein conformations requires neural networks that conserve and exclude certain geometric and symmetry properties of coordinate data. We next asked if writhe could be a useful geometric property to parameterize SE(3)-equivariant neural network architectures for use in generative models of protein structures and IDP structural ensembles. We hypothesized that writhe would be useful for parameterizing SE(3)-equivariant functions because it behaves identically to a Euclidean distance under rotations and translations, but changes sign (is equivariant) under reflections, making it a *parity-odd pseudoscalar*. As a result, scalar functions of the writhe distinguish mirror-image reflected protein conformations, while functions of Euclidean distances alone cannot. In Supplementary Figure S11, we show that the set of Euclidean distances for a structure and its mirror image are identical (invariant to parity), while the writhe for a structure and its mirror image is distinguished exactly by a change in sign (odd parity). This demonstrates that writhe can be considered equivariant to parity.

As a proof-of-principle, we leverage the odd-parity symmetry of writhe to design an efficient SE(3)-equivariant neural network to sample IDP conformations with a score-based denoising diffusion probabilistic model (DDPM)^44^. We build our model by integrating functions of writhe (see Methods, Supplementary Information Appendix B) into the previously reported, E(3)- equivariant, polarizable atom interaction network (PaiNN)^34, 41^ to obtain an SE(3)-equivariant model. The PaiNN architecture is a message passing graph neural network (MPNN) designed for molecular property prediction that has been used for protein structure generation in previous studies.^34^ The PaiNN architecture uses direction vectors between atoms and their magnitudes (Euclidean distances) to parameterize equivariant functions, making it efficient and scalable to high-dimensional MD datasets composed of many samples. Previous work has shown that the original PaiNN architecture systematically generates conformations and their mirror images with equal likelihood due to its E(3)-equivariant symmetry.^34, 40^ On account of their symmetry, E(3)- equivariant generative models sample protein structures comprised of amino acids with inverted chirality in all-atom models. Here, we observe that when this architecture is applied to generate IDP ensembles with a one-particle-per-residue resolution, popular in coarse-grained IDP models, it can also invert the chirality (i.e., writhe) of backbone chain crossings (*vide infra*, Figure 6).

**Figure 6.**
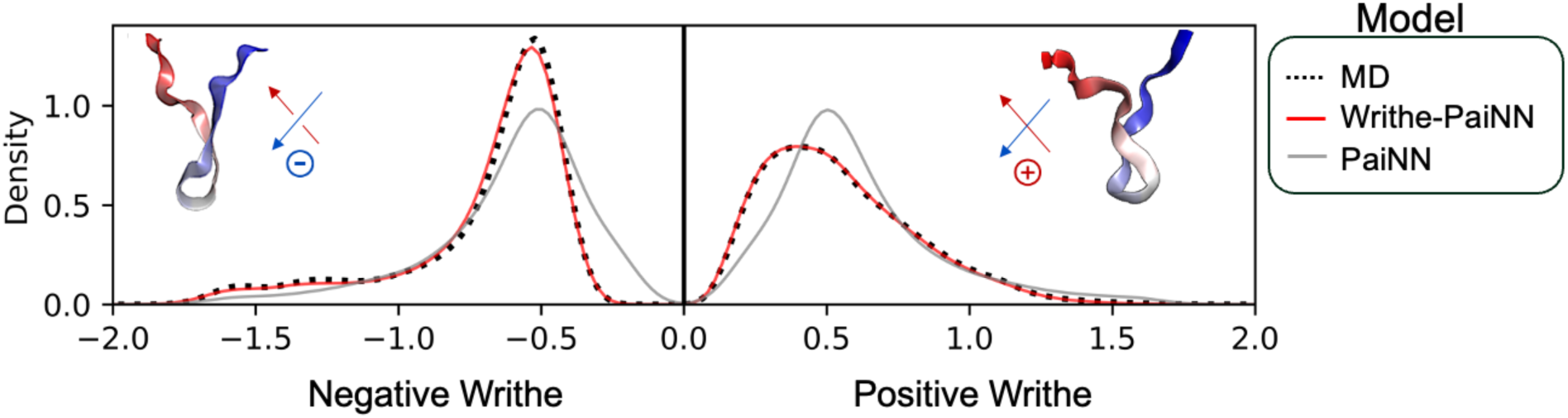
A writhe-based, SE(3) denoising diffusion probabilistic model (DDPM) accurately reproduces the populations of positive writhe and negative writhe backbone chain crossings observed in an all-atom MD simulation. We compare the populations of positive and negative Wr*_l_*_=1_ chain crossings observed in a target all-atom MD ensemble of a 20-residue C-terminal fragment of α-synuclein and ensembles obtained from DDPMs trained using the E(3)-equivariant PaiNN architecture and the SE(3)-equivariant Writhe-PaiNN architecture. To compare the relative populations of positive and negative chain crossings in each ensemble, we take separate sums of negative Wr*_l_*_=1_ values and positive Wr*_l_*_=1_ values in each frame and compare the distributions of these sums obtained from each ensemble with a kernel density estimate.

To obtain a computationally efficient model that appropriately distinguishes the populations of structures and their mirror images, we modify the PaiNN architecture using SE(3)-equivariant functions derived from writhe and cross-products between each pair of segments (Supplementary Information Appendix A).^66, 67^ To construct message passing neural network layers between atoms, we derive a writhe-graph Laplacian^68^ that maps pair-wise writhe features between segments to pair-wise writhe features between atoms (see Methods and Supplementary Information Appendix B). We refer to this neural network architecture as “Writhe-PaiNN”. As a proof of principle, we train denoising diffusion probabilistic models (DDPMs) using the original PaiNN and the Writhe-PaiNN neural network architectures on Cα-coordinate data obtained from a previously published^69^ 100 μs MD simulation of a 20-residue C-terminal fragment of the intrinsically disordered protein α-synuclein and use both DDPMs to generate Cα-coordinate ensembles of this fragment.

To demonstrate that the Writhe-PaiNN architecture appropriately models the chirality of generated structures and achieves SE(3) equivariant symmetry, we compare the populations of positive writhe and negative writhe crossings in ensembles generated by the Writhe-PaiNN and PaiNN architectures (Figure 6). To compare the relative populations of positive and negative crossings, we separately sum the negative Wr*_l_*_=1_ values and positive Wr*_l_*_=1_ values in each frame and compare the distributions of these sums obtained from each ensemble with a kernel density estimate. We observe a clear asymmetry in the distribution of positive and negative Wr*_l_*_=1_ values in the target MD ensemble training data, with a substantially larger population of negative writhe crossings (Figure 6). We observe that the distributions of negative writhe crossings and positive writhe crossings obtained from the DDPM trained with the E(3) PaiNN architecture are symmetric, in disagreement with the original MD trajectory. In contrast, the DDPM trained using the SE(3)- equivariant Writhe-PaiNN architecture accurately reproduces the populations of positive and negative crossings observed in the original MD trajectory.

For parity-invariant observables like the radius of gyration and an intramolecular bend-angle formed by Cα atoms 1, 10, and 20, we observe that the distributions obtained from the PaiNN and Writhe-PaiNN DDPMs are in close agreement, indicating that Writhe-PaiNN architecture only impacts parity equivariance (Supplementary Figure S12). These results demonstrate that augmenting the E(3) symmetry of the PaiNN architecture to SE(3) using our writhe-based approach yields an efficient SE(3)-equivariant neural network architecture that accurately reproduces differences in the chirality of chain crossings observed in MD simulation coordinates.^40, 34^ Further discussion of our implementation of the PaiNN architecture and model training protocols is provided in Supplementary Information Appendix C, “PaiNN architecture implementation and DDPM training”.

## Conclusion

Our results demonstrate that writhe-based structural descriptors provide a powerful basis to capture slow dynamic processes, metastable states, and large-scale conformational transitions in IDPs. By leveraging the geometric and topological properties of writhe, we develop a multiscale description of IDP ensembles that identifies kinetically distinct conformational states more effectively than traditional distance-based features. We find that writhe describes slow conformational changes in IDPs processes more effectively than Euclidean distances because it changes sign (is equivariant) under mirror reflection and therefore distinguishes the chirality of local and global structural features of IDPs that are not distinguished by Euclidean distances. We show that multiscale writhe descriptors provide a general and robust framework to describe structural and kinetic ensembles of IDPs by applying these descriptors to analyze long-timescale MD simulations of a diverse set of IDPs and a fast-folding protein. We demonstrate that writhe features consistently outperform Euclidean distances in describing the kinetic variance of MD trajectories and facilitate the construction of Markov state models (MSMs) that describe longer timescale dynamics. These findings highlight the potential of using writhe as a general framework for analyzing high-dimensional conformational landscapes of IDPs.

We further demonstrate that the symmetry properties of writhe can be used to build an SE(3)- equivariant neural network architecture and that this architecture can be used to construct a generative model of an IDP ensemble. Specifically, we incorporate writhe into the PaiNN neural network architecture, augmenting its symmetry from E(3) to SE(3). We apply this framework to train a denoising diffusion probabilistic model (DDPM) on an IDP conformational ensemble from a long-timescale all-atom MD simulation. Our results demonstrate that the generated conformational ensemble accurately reproduces the MD distribution of chiral backbone chain crossings, while the distribution obtained from a DDPM trained with an E(3)-equivariant network fails to differentiate chiral crossings.

We emphasize that the DDPMs presented in this work are trained on a single MD simulation dataset to evaluate the ability of each neural network architecture to model the symmetry of chiral chain crossings and faithfully reproduce a target ensemble. Our generative modelling results are presented as a proof-of-principle to illustrate that the symmetry properties of writhe can be exploited to parameterize SE(3) equivariant neural networks for protein structures. Scaling our model and training data to generalize to arbitrary IDP sequences will be explored in future work.

Our findings demonstrate that writhe-based descriptors can be applied to improve the resolution of structural and kinetic models of IDPs and data-driven approaches for modeling IDP ensembles. In addition to improving the quality of MSMs and providing a new tool to incorporate into training generative models, we anticipate that writhe descriptors may be valuable for evaluating and improving enhanced sampling approaches for IDP simulations. As writhe is a slowly varying order parameter in MD simulations of IDPs, it may serve as an effective collective variable for biasing enhanced sampling all-atom MD simulations, such as metadynamics^70^ or umbrella sampling^71^, to efficiently explore rare conformational transitions in IDPs. Extensions of writhe-based approaches, including using higher-order writhe descriptors such as those described by Rogen and coworkers,^30, 48^ may also be useful for developing improved dimensionality reduction and clustering methods for IDPs. Writhe descriptors could potentially serve as global shape coordinates for applications in autoencoders and VAMPnets^64^, facilitating interpretable representations of IDP state spaces. We anticipate that the writhe descriptors described here may be valuable for assessing the topological complexity of coarse-grained IDP models and generative models of IDPs, to identify areas where those models can be improved to more closely model ensembles obtained from all-atom MD.

Our results demonstrate that writhe is a powerful descriptor of IDP conformational ensembles, capable of enhancing the analysis of molecular simulations and improving machine learning approaches for understanding the behavior of intrinsically disordered proteins (IDPs). To facilitate the use of writhe to analyze protein ensembles, we provide an open-source Python package for computing writhe-based descriptors, which we anticipate will serve as a valuable resource for the structural biology and biophysics communities.

## Methods

### Markov state models and time-lagged canonical correlation analysis (tCCA)

In the context of protein biophysics, Markov state models (MSMs) are multiscale stochastic models used to describe the dynamics of transitions between discrete conformational states.^59, 72^ Under the assumption that interconversions between states are approximately Markovian, MSMs are a rigorous tool to predict dynamic and stationary experimental observables from MD simulation data. MSMs are validated through self-consistency measures. Physical observables like relaxation timescales predicted by MSMs should be invariant to the model’s lag time, and the evolution of the transition matrix should adhere to the Chapman-Kolmogorov equation.^72–74^ However, the practical utility of MSMs and the insight they provide is based on their spatiotemporal resolution. Loosely speaking, optimizing the spatial-temporal resolution of a model is equivalent to finding the model that is valid at the shortest lag time and has the largest number of kinetically distinct states whose transition statistics are sampled sufficiently.

To visualize feature sets and provide a numerical quantification of their usefulness in constructing kinetic models, we utilize time-lagged canonical correlation analysis (tCCA).^15, 16, 20^ Canonical correlation analysis (CCA) can be viewed as a dimensionality reduction that relates two sets of variables (or datasets) by finding orthogonal transformations (or linear combinations) of each that maximize their correlation.^21, 22^ CCA is computed by the singular value decomposition of the whitened correlation matrix of the two datasets. Here, we consider two datasets composed of the same number of *n* samples and *d* features, X and Y ∈ ℝ^𝑛×𝑑^and the averages of each of the *d* features, 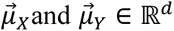. The CCA decomposition can be written in terms of sample covariance matrices (𝐶_∗_):

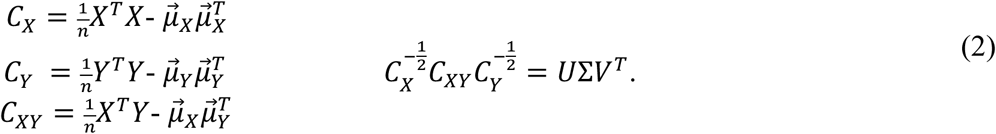

Here, *U* and *V* are the left and right singular functions, respectively, that yield the orthogonal transformations and Σ is a matrix with the singular values (𝜎_𝑖_) on the diagonal and zeros everywhere else. The singular values are the correlations of the transformed data (see reference 17 for more details). Low-dimensional representations of the data are obtained by projecting it onto the dominant singular functions, i.e., those with the largest singular values.

Time-lagged canonical correlation analysis is a special case of CCA where the datasets are time-lagged versions of each other. Therefore, tCCA finds projections of the data with maximal *autocorrelation*. In this case, the singular values are the *autocorrelations* of the transformed data. If the data is sampled from equilibrium, the sum of the squared singular values describes the *kinetic variance* captured by their corresponding singular functions and can be used as a variational score (or VAMP-2 score)^19^ to find an optimal set of input features for capturing slow processes and building MSMs.^15–20^ The kinetic variance is defined as:

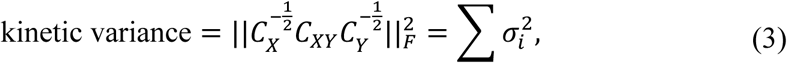

where 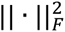 denotes the Frobenius norm of the whitened correlation matrix. While this exact expression is sometimes defined as a VAMP-2 score, we use the term *kinetic variance* to differentiate from the contexts where a VAMP-2 score is used as an optimization target for VAMPnets^64^ and related deep learning approaches for constructing MSMs.^6, 63^

tCCA is closely related to time-independent component analysis (tICA).^11, 22, 75^ Both find projections of the data with maximal autocorrelation, however, tCCA is more general as it can natively handle off-equilibrium statistics due to its formulation using the SVD. In contrast, tICA explicitly enforces reversibility by symmetrizing the autocovariance matrix between the instantaneous (A) and time-lagged data (B) to obtain real eigenvalues (𝚲) and orthonormal eigenvectors (𝑉) from the generalized eigenvalue problem: 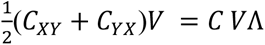 where 𝑋 and 𝑌 are time-lagged versions of the same dataset and the sample covariance matrix of the full dataset is denoted as the matrix, 𝐶 .

### Markov State Model Construction

For all systems, we build MSMs by first computing writhe and Euclidean distance features. We determine the best writhe feature set for each system by performing tCCA on several combinations of writhe features and computing the kinetic variance of each projection, as shown in Supplementary Figure S1. The writhe feature set with the largest kinetic variance score is considered optimal and utilized in further analysis. We proceed by building two MSMs for each system – one using the optimal writhe feature set and the other using inter-residue distances for comparison. In either case, projections of the features onto a variable number of tCCA components (2, 3, 5, and 10) are used to cluster the trajectory over a range of K-means clusters (10, 20, and 40). The clusters obtained from all combinations of tCCA components and K-means clusters are utilized to estimate MSMs over a range of lag times. We scan MSM results by plotting the longest implied timescale (ITS) from each model as a function of the lag time (Supplementary Figure S10). From this, we determine the suitable combinations of tCCA dimensions, K-means clusters, and model lag time.

We identified suitable hyperparameters for the MSMs of Aβ42 and ACTR using the grid search defined above, and proceeded to construct coarse-grained models with a small number of states. To construct coarse-grained models, we found that using 40 initial clusters and 3 tCCA components strikes the best balance between interpretable and reproducible metastable state definitions and capturing slow dynamical processes. For Aβ42, we used an MSM lag time of τ = 2.5 ns (shortest lag time with converged ITS) to increase statistical efficiency, given that the simulation data is comprised of thousands of short trajectories (maximum length ∼ 90 ns). For ACTR, we utilized a MSM lag time of τ = 6 ns based on ITS convergence and generalization at longer lag times (Supplementary Figure S8). We determined the number of metastable states for each model based on the number of ITS resolved by the MSM and the consistency of the PCCA++ algorithm in identifying the same set of metastable states from an ensemble of bootstrapped MSMs. All MSM observables are reported with 95% confidence intervals obtained from bootstrapped ensembles of MSMs containing 1000 samples generated using *Bayesian Markov models*.^72, 73^ Mean first passage times (MFPTs) and transition probability fluxes were computed using transition path theory^76–78^ analysis. MSM estimation and transition path theory analysis were performed using the *deeptime*^79^ Python software package.

### Score-based generative models

Score-based generative diffusion models are probabilistic generative models used to infer independent samples from a data distribution by learning a so-called *score field* that reverses (or denoises) a time-inhomogeneous stochastic process that gradually corrupts data to random noise.^44, 80–83^ The data distribution, 𝑝(𝑥^0^), is gradually transformed to a simple prior distribution, 𝑝(𝑥^𝑇^), through the following stochastic differential equation (SDE) in Ito form^44, 82^:

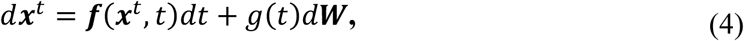

where t is a continuous time variable defined over [0, T] referred to as the diffusion time, W is the standard Wiener process, 𝒇(·, 𝑡) is a known vector valued function referred to as the drift coefficient and 𝑔(·) is often treated as (and is here) a known scalar function referred to as the diffusion coefficient of 𝑥(𝑡). A *backward diffusion* process described by the following is used to transform samples from a simple prior, 𝑝(𝑥^𝑇^), to samples from the data distribution.^44, 84^

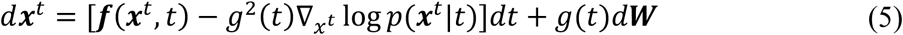

Where 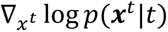 is the score field and can be approximated by a deep neural network that directs samples from a simple prior to the data distribution via a series of noisy perturbations in the direction of maximum likelihood.

#### Geometric deep learning

Here, we define a function, 𝑓, as ‘invariant’ under a group-action 𝑔 if 𝑓(𝑥) = 𝑓(𝑆_g_𝑥) and ‘equivariant’ if 𝑇_g_𝑓(𝑥)= 𝑓(𝑆_g_𝑥), where 𝑆_g_ and 𝑇_g_ are linear representations of the group element 𝑔.^34^ For molecular coordinates free to globally translate and rotate in 3 dimensions, the relevant symmetry groups are the *special Euclidean group* (proper rotations and translations), SE(3), and the *Euclidean group* (proper rotations, translations, and parity or reflections), E(3). It has been shown that probability distributions estimated from score-based diffusion models are invariant to the transformations their corresponding score fields are equivariant to.^40^ This can be leveraged to guide the construction of models using physical principles. For chiral molecules like proteins sampled from MD, predicted distributions should be invariant to rotations and translations because these transformations do not change the conformational state of the molecule. Thus, we require an SE(3)-equivariant neural network.

Here, we construct an SE(3)-equivariant model by modifying the symmetry of the E(3) equivariant polarizable atom interaction neural network (PaiNN).^41^ We modify the PaiNN architecture to align with the general formulism of SE(3)-equivariant vector functions based on invariant scalars given by Villar et. al.^66, 67^ PaiNN is a message passing graph neural network architecture that parameterizes equivariant functions using invariant scalar features (𝑠_𝑖_), equivariant vectorial features 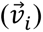, inter-atom direction vectors 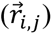, and Euclidean distances, 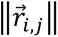. Here, *i* and *j* index atoms of a molecular structure. All features used in the model are obtained from atomic coordinates, apart from the invariant scalar features (𝑠_𝑖_) and equivariant vectorial features 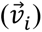 of each atom, which are used internally to govern the symmetry properties of the model and make predictions. Invariant scalars (𝑠_𝑖_) are updated in the message block of PaiNN using atom wise continuous filter convolutions^85^ parameterized by Euclidean distances and invariant scalar features: 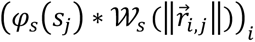, where 𝜑 denotes a generic multilayer perceptron (MLP) and W is an MLP composed with a cosine (+) and sine (-) positional encoding: 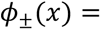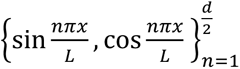, that embeds distances, 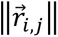, to the dimension of the model.^41^ We use a similar approach for scalar writhe features, except positional encodings of writhe features consist of only sines to retain their odd parity. We denote sine only positional encodings as: 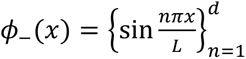. We incorporate atom-wise scalar writhe features (Appendix A), 𝑤_i,j_, into continuous filter convolutions by concatenating embedded scalar writhe features and Euclidean distances. We denote the concatenated scalar writhe and distance features as, 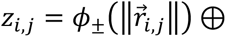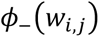, where ⊕ denotes concatenation across the feature dimension. The residual of the scalar message (*m*) update function is defined as:

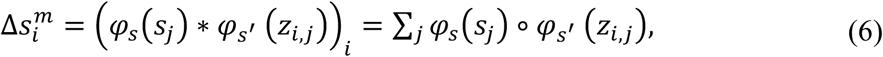

where the sum is taken over the *j* neighbors of atom *i* to update its invariant scalar features, 𝑠_𝑖_, and o denotes the Hadamard product. We modify the equivariant vector function of PaiNN to SE(3) using the cross-product vector between segments (𝑻_1_ × 𝑻_2_ in Supplementary Figure S1) obtained from the computation of each scalar value of the writhe, 𝑤_𝑖,j_. In the following, we denote the writhe-derived cross-product vectors as 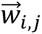 for ease of notation. Similarly to our treatment of the scalar writhe and Euclidean distances, we incorporate cross product vectors 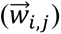 following the same approach as the inter-atom direction vectors 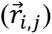 in the original PaiNN architecture by including 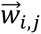 in the weighted sum of equivariant vectors used to update equivariant vector features, 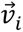. The residual of the vector message (*m*) update function is defined as:

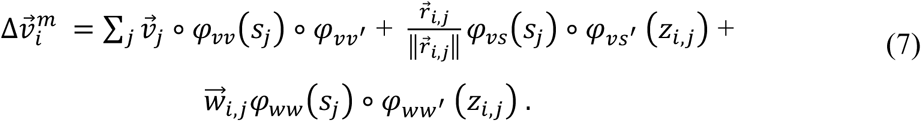

## Author Contributions

PR conceived, designed, and supervised the research. PR and TRS conceived and designed the trajectory analysis and dynamic modeling portion of the research. TRS and SO conceived and designed the generative deep learning portion of the paper while TRS was visiting Chalmers University of Technology. TRS and PR wrote the paper. TRS, PR and SO edited and revised the paper.

## Acknowledgements

The authors acknowledge Peter Røgen for valuable discussions and for sharing code, and Mathias Schreiner for valuable discussions regarding the implementation of writhe in the PaiNN neural network architecture.

## Data and Code Availability

All code for reproducing the MD trajectory analyses in this paper is freely available from GitHub (https://github.com/paulrobustelli/Sisk_IDP_writhe_2025). A general-purpose implementation of the methods developed in this study for computing writhe and analyzing molecular dynamics simulation data is available as the open-source Python package `writhe_tools`, which is freely distributed via the Python Package Index (PyPI) and can be installed using `pip install writhe_tools.` The freely available MD trajectories of a-synuclein, ACTR, PaaA2, HP35, and N_TAIL_ analyzed in this work are available for non-commercial use by request from D.E. Shaw Research. MD trajectories of Ab42 are freely available from https://zenodo.org/record/4247321.

## References

(1) Babu, M. M.; van der Lee, R.; de Groot, N. S.; Gsponer, J. Intrinsically disordered proteins: regulation and disease. Current opinion in structural biology 2011, 21 (3), 432–440.

(2) Robustelli, P.; Piana, S.; Shaw, D. E. Developing a molecular dynamics force field for both folded and disordered protein states. Proceedings of the National Academy of Sciences 2018, 115 (21), E4758–E4766.

(3) Piana, S.; Robustelli, P.; Tan, D.; Chen, S.; Shaw, D. E. Development of a Force Field for the Simulation of Single-Chain Proteins and Protein–Protein Complexes. Journal of Chemical Theory and Computation 2020, 16 (4), 2494–2507.

(4) Borthakur, K.; Sisk, T. R.; Panei, F. P.; Bonomi, M.; Robustelli, P. Determining accurate conformational ensembles of intrinsically disordered proteins at atomic resolution. bioRxiv 2024.

(5) Best, R. B.; Zheng, W.; Mittal, J. Balanced Protein–Water Interactions Improve Properties of Disordered Proteins and Non-Specific Protein Association. Journal of Chemical Theory and Computation 2014, 10 (11), 5113–5124.

(6) Sisk, T. R.; Robustelli, P. Folding-upon-binding pathways of an intrinsically disordered protein from a deep Markov state model. Proceedings of the National Academy of Sciences 2024, 121 (6).

(7) Löhr, T.; Kohlhoff, K.; Heller, G. T.; Camilloni, C.; Vendruscolo, M. A kinetic ensemble of the Alzheimer’s Aβ peptide. Nature Computational Science 2021, 1 (1), 71–78.

(8) Bonomi, M.; Heller, G. T.; Camilloni, C.; Vendruscolo, M. Principles of protein structural ensemble determination. Current Opinion in Structural Biology 2017, 42, 106–116.

(9) Lindorff-Larsen, K.; Trbovic, N.; Maragakis, P.; Piana, S.; Shaw, D. E. Structure and Dynamics of an Unfolded Protein Examined by Molecular Dynamics Simulation. Journal of the American Chemical Society 2012, 134 (8), 3787–3791.

(10) Coyle, D.; Hampton, L. 21st century progress in computing. Telecommunications Policy 2023, 48 (1), 102649–102649.

(11) Noé, F.; Clementi, C. Kinetic Distance and Kinetic Maps from Molecular Dynamics Simulation. Journal of Chemical Theory and Computation 2015, 11 (10), 5002–5011.

(12) Brunton, S. L.; Proctor, J. L.; Kutz, J. N. Discovering governing equations from data by sparse identification of nonlinear dynamical systems. Proceedings of the National Academy of Sciences 2016, 113 (15), 3932–3937.

(13) Lu, H.; Tartakovsky, D. M. Extended dynamic mode decomposition for inhomogeneous problems. Journal of Computational Physics 2021, 444, 110550.

(14) Husic, B. E.; Pande, V. S. Markov State Models: From an Art to a Science. Journal of the American Chemical Society 2018, 140 (7), 2386–2396.

(15) Wu, H.; Noé, F. Variational Approach for Learning Markov Processes from Time Series Data. Journal of Nonlinear Science 2019, 30 (1), 23–66.

(16) Noé, F.; Nüske, F. A Variational Approach to Modeling Slow Processes in Stochastic Dynamical Systems. Multiscale Modeling & Simulation 2013, 11 (2), 635–655.

(17) Nüske, F.; Keller, B. G.; Pérez-Hernández, G.; Mey, A. S. J. S.; Noé, F. Variational Approach to Molecular Kinetics. Journal of Chemical Theory and Computation 2014, 10 (4), 1739–1752.

(18) McGibbon, R. T.; Pande, V. S. Variational cross-validation of slow dynamical modes in molecular kinetics. The Journal of Chemical Physics 2015, 142 (12), 124105.

(19) Wu, H.; Nüske, F.; Paul, F.; Klus, S.; Koltai, P.; Noé, F. Variational Koopman models: Slow collective variables and molecular kinetics from short off-equilibrium simulations. The Journal of Chemical Physics 2017, 146 (15), 154104.

(20) Scherer, M. K.; Husic, B. E.; Hoffmann, M.; Paul, F.; Wu, H.; Noé, F. Variational selection of features for molecular kinetics. The Journal of Chemical Physics 2019, 150 (19), 194108.

(21) Hotelling, H. Relations Between Two Sets of Variates. Biometrika 1936, 28 (3/4), 321.

(22) Molgedey, L.; Schuster, H. G. Separation of a mixture of independent signals using time delayed correlations. Physical Review Letters 1994, 72 (23), 3634–3637.

(23) Călugăreanu, G. L’intégrale de Gauss et l’analyse des nœuds tridimensionnels. Rev. Math. Pures Appl. 1959, *4*, 5–20.

(24) Fuller, F. B. The Writhing Number of a Space Curve. Proceedings of the National Academy of Sciences 1971, 68 (4), 815–819.

(25) Rackovsky, S.; Scheraga, H. A. Differential Geometry and Polymer Conformation. 1. Comparison of Protein Conformations. Macromolecules 1978, 11 (6), 1168–1174.

(26) Rackovsky, S.; Scheraga, H. A. Differential Geometry and Polymer Conformation. 2. Development of a Conformational Distance Function. Macromolecules 1980, 13 (6), 1440–1453.

(27) Zhi, D.; Shatsky, M.; Brenner, S. E. Alignment-free local structural search by writhe decomposition. Bioinformatics 2010, 26 (9), 1176–1184.

(28) Konstantin, K.; Langowski, J. Computation of writhe in modeling of supercoiled DNA. Biopolymers 2000, 54 (5), 307–317.

(29) Levitt, M.; Lifson, S. Refinement of protein conformations using a macromolecular energy minimization procedure. Journal of Molecular Biology 1969, 46 (2), 269–279.

(30) Røgen, P.; Bohr, H. A new family of global protein shape descriptors. Mathematical Biosciences 2003, 182 (2), 167–181.

(31) Røgen, P.; Karlsson, P. W. Parabolic section and distance excess of space curves applied to protein structure classification. Geometriae Dedicata 2008, 134 (1), 91–107.

(32) Janson, G.; Feig, M. Transferable deep generative modeling of intrinsically disordered protein conformations. PLOS Computational Biology 2024, 20 (5), e1012144.

(33) Gupta, A.; Dey, S.; Hicks, A.; Zhou, H. X. Artificial intelligence guided conformational mining of intrinsically disordered proteins. Communications biology 2022, 5 (1).

(34) Schreiner, M.; Winther, O.; Olsson, S. Implicit Transfer Operator Learning: Multiple Time-Resolution Surrogates for Molecular Dynamics. In 37th Conference on Neural Information Processing Systems, 2023.

(35) Noé, F.; Olsson, S.; Köhler, J.; Wu, H. Boltzmann Generators: Sampling Equilibrium States of Many-Body Systems with Deep Learning. Science 2019, 365 (6457), eaaw1147.

(36) Jumper, J.; Evans, R.; Pritzel, A.; Green, T.; Figurnov, M.; Ronneberger, O.; Tunyasuvunakool, K.; Bates, R.; Žídek, A.; Potapenko, A.;, et al. Highly Accurate Protein Structure Prediction with Alphafold. Nature 2021, 596 (7873), 583–589.

(37) Abramson, J.; Adler, J.; Dunger, J.; Evans, R.; Green, T.; Pritzel, A.; Ronneberger, O.; Willmore, L.; Ballard, A. J.; Bambrick, J.;, et al. Accurate structure prediction of biomolecular interactions with AlphaFold 3. Nature 2024, 630 (630), 493–500.

(38) Lewis, S.; Hempel, T.; Jiménez-Luna, J.; Gastegger, M.; Xie, Y.; Foong, A. Y. K.; García Satorras, V.; Abdin, O.; Veeling, B. S.; Zaporozhets, I.; et al. Scalable Emulation of Protein Equilibrium Ensembles with Generative Deep Learning. *bioRxiv* 2024.

(39) Novak, A.; Lotthammer, J. M.; Emenecker, R. J.; Holehouse, A. S. Accurate predictions of conformational ensembles of disordered proteins with STARLING. bioRxiv 2025.

(40) Köhler, J.; Klein, L.; Noe, F. Equivariant Flows: exact likelihood generative learning for symmetric densities. International Conference on Machine Learning 2020, 1, 5361–5370.

(41) Schütt, K.; Unke, O.; Gastegger, M. Equivariant Message Passing for the Prediction of Tensorial Properties and Molecular Spectra; https://proceedings.mlr.press/v139/schutt21a/schutt21a.pdf.

(42) Tesei, G.; Trolle, A. I.; Jonsson, N.; Betz, J.; Knudsen, F. E.; Pesce, F.; Johansson, K. E.; Lindorff-Larsen, K. Conformational ensembles of the human intrinsically disordered proteome. Nature 2024, 626, 1–8.

(43) Tesei, G.; Lindorff-Larsen, K. Improved predictions of phase behaviour of intrinsically disordered proteins by tuning the interaction range. Open Research Europe 2022, 2, 94.

(44) Song, Y.; Jascha, S.-D.; Kingma, D. P.; Kumar, A.; Stefano, E.; Poole, B. Score-Based Generative Modeling through Stochastic Differential Equations. International Conference on Learning Representations 2021.

(45) Kauffman, L. H. Formal knot theory; Dover Publications, 2006.

(46) Kauffman, L. H. Knots and Physics; World Scientific, 1994.

(47) Røgen, P.; Fain, B. Automatic classification of protein structure by using Gauss integrals. Proceedings of the National Academy of Sciences 2002, 100 (1), 119–124.

(48) Røgen, P.; Sinclair, R. Computing a New Family of Shape Descriptors for Protein Structures. Journal of Chemical Information and Computer Sciences 2003, 43 (6), 1740–1747.

(49) Bar-Natan, D. On the Vassiliev knot invariants. Topology 1995, 34 (2), 423–472.

(50) Piana, S.; Lindorff-Larsen, K.; Shaw, D. E. Protein folding kinetics and thermodynamics from atomistic simulation. Proceedings of the National Academy of Sciences 2012, 109 (44), 17845–17850.

(51) Pietrucci, F.; Laio, A. A Collective Variable for the Efficient Exploration of Protein Beta-Sheet Structures: Application to SH3 and GB1. Journal of Chemical Theory and Computation 2009, 5 (9), 2197–2201.

(52) Robustelli, P.; Piana, S.; Shaw, D. E. Mechanism of Coupled Folding-upon-Binding of an Intrinsically Disordered Protein. Journal of the American Chemical Society 2020, 142 (25), 11092–11101.

(53) Best, R. B.; Hummer, G. Optimized Molecular Dynamics Force Fields Applied to the Helix−Coil Transition of Polypeptides. Journal of Physical Chemistry B 2009, 113 (26), 9004–9015.

(54) Lindorff-Larsen, K.; Piana, S.; Palmo, K.; Maragakis, P.; Klepeis, J. L.; Dror, R. O.; Shaw, D. E. Improved side-chain torsion potentials for the Amber ff99SB protein force field. Proteins: Structure, Function, and Bioinformatics 2010, 78 (8), NA-NA.

(55) MacKerell, A. D.; Bashford, D.; Bellott, M.; Dunbrack, R. L.; Evanseck, J. D.; Field, M. J.; Fischer, S.; Gao, J.; Guo, H.; Ha, S.;, et al. All-Atom Empirical Potential for Molecular Modeling and Dynamics Studies of Proteins†. The Journal of Physical Chemistry B 1998, 102 (18), 3586–3616.

(56) Piana, S.; Lindorff-Larsen, K.; Shaw, David E. How Robust Are Protein Folding Simulations with Respect to Force Field Parameterization? Biophysical Journal 2011, *100* (9), L47-L49.

(57) MacQueen, J. Some Methods for Classification and Analysis of Multivariate Observations. In Proceedings of the Fifth Berkeley Symposium on Mathematical Statistics and Probability, Volume 1: Statistics, Berkeley, CA, USA; 1967.

(58) Rousseeuw, P. J. Silhouettes: a Graphical Aid to the Interpretation and Validation of Cluster Analysis. Journal of Computational and Applied Mathematics 1987, 20 (0377-0427), 53–65.

(59) Trendelkamp-Schroer, B.; Wu, H.; Paul, F.; Noé, F. Estimation and uncertainty of reversible Markov models. The Journal of Chemical Physics 2015, 143 (17), 174101.

(60) Röblitz, S.; Weber, M. Fuzzy spectral clustering by PCCA+: application to Markov state models and data classification. Advances in Data Analysis and Classification 2013, 7 (2), 147–179.

(61) Ziegel, J.; Röblitz, S.; Weber, M. Perron Cluster Analysis and Its Connection to Graph Partitioning for Noisy Data; Zuse Institute Berlin (ZIB), 2004. https://schlieplab.org/Static/Publications/ZR-04-39.pdf.

(62) Weber, M.; Kube, S. *Robust Perron Cluster Analysis for Various Applications in Computational Life Science*; Zuse Institute Berlin (ZIB), 2005. https://citeseerx.ist.psu.edu/document?doi=60d3416555c49096747132e68d5a9940bd19819b.

(63) Mardt, A.; Pasquali, L.; Noé, F.; Wu, H. Deep learning Markov and Koopman models with physical constraints. Proceedings of Machine Learning Research 2020, 107, 451–475.

(64) Mardt, A.; Pasquali, L.; Wu, H.; Noé, F. VAMPnets for deep learning of molecular kinetics. Nature Communications 2018, 9 (1), 1–11.

(65) Monteiro da Silva, G.; Cui, J. Y.; Dalgarno, D. C.; Lisi, G. P.; Rubenstein, B. M. High-throughput prediction of protein conformational distributions with subsampled AlphaFold2. Nature Communications 2024, 15 (1), 2464.

(66) Blum-Smith, B.; Villar, S. Machine Learning and Invariant Theory. Notices of the American Mathematical Society 2023, 70 (08), 1–1.

(67) Chen, N.; Villar, S. SE(3)-equivariant self-attention via invariant features; 2022. https://ml4physicalsciences.github.io/2022/files/NeurIPS_ML4PS_2022_154.pdf.

(68) Strang, G. Linear algebra and learning from data; Wellesley-Cambridge Press, 2019.

(69) Robustelli, P.; Ibanez-de-Opakua, A.; Campbell-Bezat, C.; Giordanetto, F.; Becker, S.; Zweckstetter, M.; Pan, A. C.; Shaw, D. E. Molecular Basis of Small-Molecule Binding to α-Synuclein. Journal of the American Chemical Society 2022, 144 (6), 2501–2510.

(70) Laio, A.; Parrinello, M. Escaping free-energy minima. Proceedings of the National Academy of Sciences of the United States of America 2002, 99 (20), 12562–12566.

(71) Torrie, G. M.; Valleau, J. P. Nonphysical sampling distributions in Monte Carlo free-energy estimation: Umbrella sampling. Journal of Computational Physics 1977, 23 (2), 187–199.

(72) Prinz, J.-H.; Wu, H.; Sarich, M.; Keller, B.; Senne, M.; Held, M.; Chodera, J. D.; Schütte, C.; Noé, F. Markov models of molecular kinetics: Generation and validation. The Journal of Chemical Physics 2011, 134 (17), 174105.

(73) Noé, F.; Rosta, E. Markov Models of Molecular Kinetics. The Journal of Chemical Physics 2019, 151 (19), 190401.

(74) Scherer, M. K.; Trendelkamp-Schroer, B.; Paul, F.; Pérez-Hernández, G.; Hoffmann, M.; Plattner, N.; Wehmeyer, C.; Prinz, J.-H.; Noé, F. PyEMMA 2: A Software Package for Estimation, Validation, and Analysis of Markov Models. Journal of Chemical Theory and Computation 2015, 11 (11), 5525–5542.

(75) Paul, F.; Wehmeyer, C.; Abualrous, E. T.; Wu, H.; Crabtree, M. D.; Schöneberg, J.; Clarke, J.; Freund, C.; Weikl, T. R.; Noé, F. Protein-peptide association kinetics beyond the seconds timescale from atomistic simulations. Nature Communications 2017, 8 (1).

(76) E, W.; Vanden-Eijnden, E. Towards a Theory of Transition Paths. Journal of Statistical Physics 2006, 123 (3), 503–523.

(77) Metzner, P.; Christof, S.; Vanden-Eijnden, E. Transition Path Theory for Markov Jump Processes. Multiscale Modeling and Simulation 2009, 7 (3), 1192–1219.

(78) Sturzenegger, F.; Zosel, F.; Holmstrom, E. D.; Buholzer, K. J.; Makarov, D. E.; Nettels, D.; Schuler, B. Transition path times of coupled folding and binding reveal the formation of an encounter complex. Nature Communications 2018, 9 (1).

(79) Hoffmann, M.; Scherer, M.; Hempel, T.; Mardt, A.; de Silva, B.; Husic, B. E.; Klus, S.; Wu, H.; Kutz, N.; Brunton, S. L.;, et al. Deeptime: a Python library for machine learning dynamical models from time series data. Machine Learning: Science and Technology 2021, 3 (1), 015009.

(80) Sohl-Dickstein, J.; Weiss, E.; Maheswaranathan, N.; Ganguli, S. Deep Unsupervised Learning using Nonequilibrium Thermodynamics. In Proceedings of the 32nd International Conference on Machine Learning, 2015; Vol. 37, pp 2256-2265.

(81) Ho, J.; Jain, A.; Abbeel, P. Denoising Diffusion Probabilistic Models. arXiv preprint arXiv:2006.11239 2020.

(82) Song, Y.; Durkan, C.; Murray, I.; Ermon, S. Maximum Likelihood Training of Score-Based Diffusion Models. *arXiv preprint arXiv:2101.09258* 2021.

(83) Maoutsa, D.; Reich, S.; Opper, M. Interacting Particle Solutions of Fokker–Planck Equations Through Gradient–Log–Density Estimation. Entropy 2020, 22 (8), 802.

(84) Anderson, B. D. O. Reverse-time diffusion equation models. Stochastic Processes and their Applications 1982, 12 (3), 313–326.

(85) Schütt, K. T.; Pieter-Jan, K.; Sauceda, H. E.; Chmiela, S.; Alexandre, T.; Klaus-Robert, M. SchNet: A continuous-filter convolutional neural network for modeling quantum interactions. arXiv (Cornell University) 2017.

