## Supplementary Information for "Characterizing structural and kinetic ensembles of intrinsically disordered proteins using writhe"

Paul Robustelli:

**Gaussian integrals and writhe of continuous curves.**

The writhe of continuous curves in $\mathbb{R}^3$ can be expressed as the Gaussian integral[1, 2],

$$\frac{1}{4\pi}\int_0^{L_1}\int_0^{L_2}\frac{\boldsymbol{T}(s_1)\times\boldsymbol{T}(s_2)\cdot\left(\boldsymbol{r}(s_1)-\boldsymbol{r}(s_2)\right)}{\|\boldsymbol{r}(s_1)-\boldsymbol{r}(s_2)\|^3}\,ds_1\,ds_2, \quad (1)$$

where $\boldsymbol{r}(s)$ is a vector valued function giving the position of each point along a space curve as a function of the arc length parameter $s$, which takes values in the interval [0, L] and spans length of the curve, L. The function $\boldsymbol{T}(s)$ is the unit tangent vector at $s$, $\boldsymbol{T}(s) = d\boldsymbol{r}/ds$, which describes the local direction of the curve. The parameters $s_1$ and $s_2$ are points on the space carve, and serve as integration variables, such that the double integral in (1) runs over all pairs of points. Evaluating the expression under the integral for two arbitrary points on the curve as a function of their arclengths $(s_1, s_2)$ and inserting the definitions of the cross and dot products yields:

$$\begin{aligned}&\text{(2.a)}\ \ \boldsymbol{r}(s_1)-\boldsymbol{r}(s_2)=\Delta\boldsymbol{r}\\ &\text{(2.b)}\ \ \boldsymbol{T}(s_1)\times\boldsymbol{T}(s_2) = \hat{\boldsymbol{n}}\sin\theta \quad\Longrightarrow\quad \frac{\boldsymbol{T}(s_1)\times\boldsymbol{T}(s_2)\cdot\Delta\boldsymbol{r}}{\|\Delta\boldsymbol{r}\|^3}=\frac{\sin\theta\hat{\boldsymbol{n}}\cdot\Delta\boldsymbol{r}}{\|\Delta\boldsymbol{r}\|^3}=\frac{\sin\theta\cos\phi}{\|\Delta\boldsymbol{r}\|^2}. \quad (2)\\ &\text{(2.c)}\ \ \hat{\boldsymbol{n}}\cdot\Delta\boldsymbol{r}=\|\Delta\boldsymbol{r}\|\cos\phi\end{aligned}$$

In the above, we have abbreviated the displacement vector between the two points on the curve as $\Delta\boldsymbol{r}$ (2.a). We use the typical definition of the cross product of unit vectors (2.b) to write $\boldsymbol{T}(s_1)\times\boldsymbol{T}(s_2)$ as a unit vector ($\hat{\boldsymbol{n}}$) orthogonal to the plane containing $\boldsymbol{T}(s_1)$ and $\boldsymbol{T}(s_2)$ multiplied by the sine of the angle between $\boldsymbol{T}(s_1)$ and $\boldsymbol{T}(s_2)$, which we denote as $\theta$. Additionally, we use the typical definition of the dot product (2.c) to write $\hat{\boldsymbol{n}}\cdot\Delta\boldsymbol{r}$ as the magnitude of $\Delta\boldsymbol{r}$ (distance between points on the curve) multiplied by the cosine of the angle between $\hat{\boldsymbol{n}}$ and $\Delta\boldsymbol{r}$ which we denote as $\phi$.

The final expression on the right-hand side of (1) illustrates that the magnitude of the angular component of the writhe ($\sin\theta\cos\phi$) reaches its maximum when the tangent vectors are orthogonal ($\theta = \pm\frac{\pi}{2}$) and their displacement ($\Delta\boldsymbol{r}$) aligns with the normal ($\hat{\boldsymbol{n}}$) of the tangent plane ($\phi = 0,\ \pi$). This occurs when $\boldsymbol{T}(s_1)$, $\boldsymbol{T}(s_2)$ and $\Delta\boldsymbol{r}$ are mutually orthogonal. We also see that the radial component ($1/\|\Delta\boldsymbol{r}\|^2$) causes the writhe to decay quadratically with the Euclidean distance between crossings. The full Gauss integral can be interpreted as the average number of signed crossings taken over all possible viewing angles in 3-space.[1, 3] The integral in equation 1 of the main text can also be written as a function of two curves to define their *Gaussian linking number*.[2, 4]

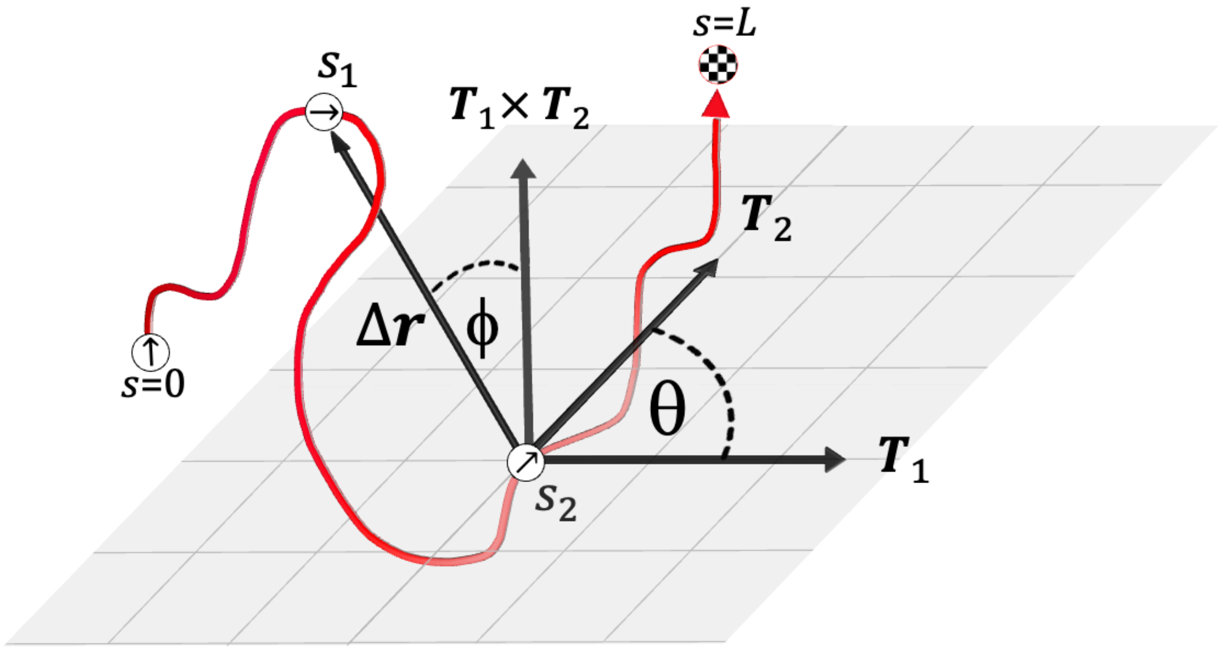


**Supplementary Figure S1: The writhe of a pair of points on a continuous space curve**. We illustrate the vectors and angles that characterize the writhe for two points ($s_1$ and $s_2$) on a continuous curve parameterized by the arclength, $s$. Here, we abbreviate the tangents at $s_1$ and $s_2$ as $\boldsymbol{T}_1$ and $\boldsymbol{T}_2$ and label the normal vector to the plane containing both tangents as their cross product, $\boldsymbol{T}_1 \times \boldsymbol{T}_2$. The direction of the tangents to the curve at $s = 0$, $s_1$, and $s_2$ are also shown as small black arrows inside white circles denoting their position on the curve. The end of the curve is labeled as $s = L$, where $L$ denotes the length of the curve. In this diagram, angle θ between the tangent vectors, $\boldsymbol{T}_1$ and $\boldsymbol{T}_2$, is $\frac{\pi}{2}$ radians or 90°, which maximizes its contribution to the angular component of the writhe. For the above diagram, the writhe monotonically increases as $\phi \to 0°$ and $\|\Delta \boldsymbol{r}\| \to 0$, i.e., as the points on the curve come close and $\Delta \boldsymbol{r}$ aligns with $\boldsymbol{T}_1 \times \boldsymbol{T}_2$, the tangent vectors appear they're crossing over a larger range of viewing angles or perspectives.

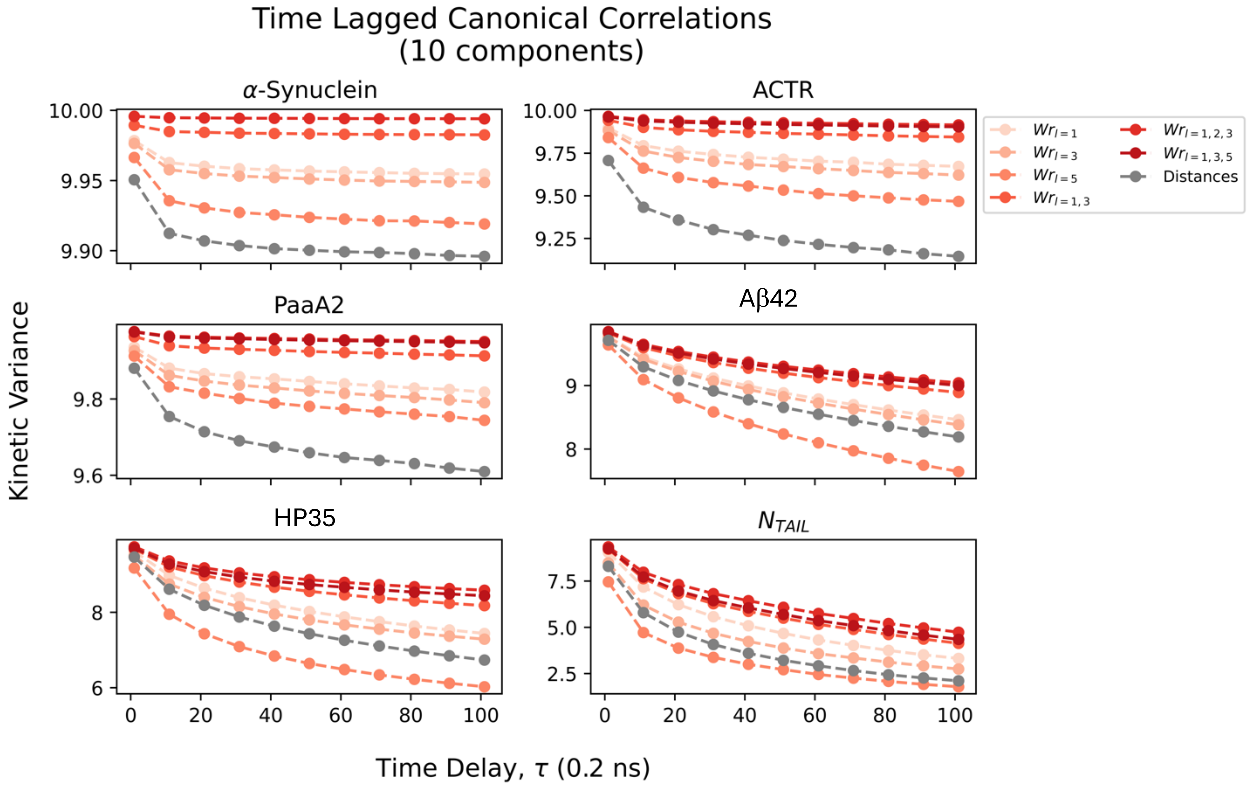


**Supplementary Figure S2: Comparison the kinetic variances (VAMP-2 scores) of writhe and Euclidian distance features from 6 long-timescale molecular dynamics simulations.** Kinetic variances obtained from performing time-lagged canonical correlation analysis[5, 6] (tCCA) for a collection of writhe feature sets (reds) and inter-residue Euclidean distances (grey) from long-timescale molecular dynamics simulations of full-length α-synuclein (140 residues), ACTR (71 residues), PaaA2 (71 residues), Aβ42 (42 residues), HP35 (35 residues) and $N_{TAIL}$ (21 residues). Here, we compute the kinetic variance from the first 10 tCCA components, making the highest possible score 10. Kinetic variances of the writhe feature set $Wr_{l=1,3,5}$ for full-length α-synuclein are not shown as we found that the high dimensionality of the dataset (~ 27,000 features) caused statical and numerical error leading to spurious results.

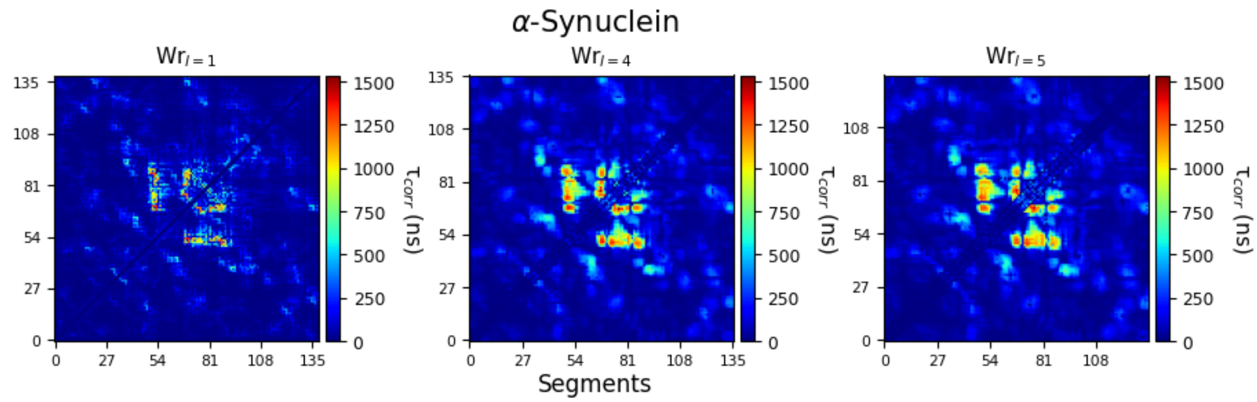


**Supplementary Figure S3-A.** Autocorrelation time ($\tau_{corr}$) matrices for pairwise writhe values between segments, computed at segment lengths of 1, 3 and 5 for a continuous long-timescale molecular dynamics simulation of α-synuclein. The autocorrelation times presented here are estimated from integrating the individual autocorrelation functions of each writhe feature up to the time that they first cross zero.

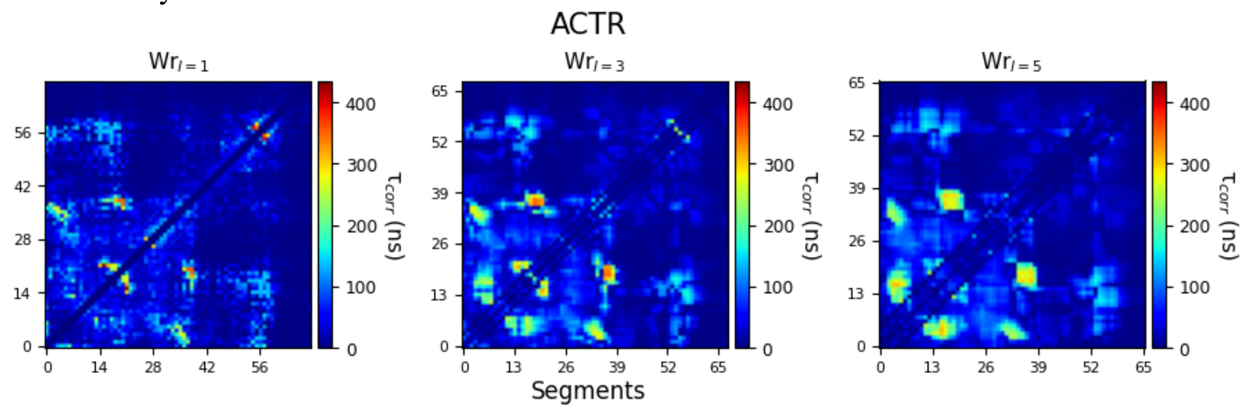


**Supplementary Figure S3-B.** Autocorrelation time ($\tau_{corr}$) matrices for pairwise writhe values between segments, computed at segment lengths of 1, 3 and 5 for a continuous long-timescale molecular dynamics simulation of ACTR. The autocorrelation times presented here are estimated from integrating the individual autocorrelation functions of each writhe feature up to the time that they first cross zero.

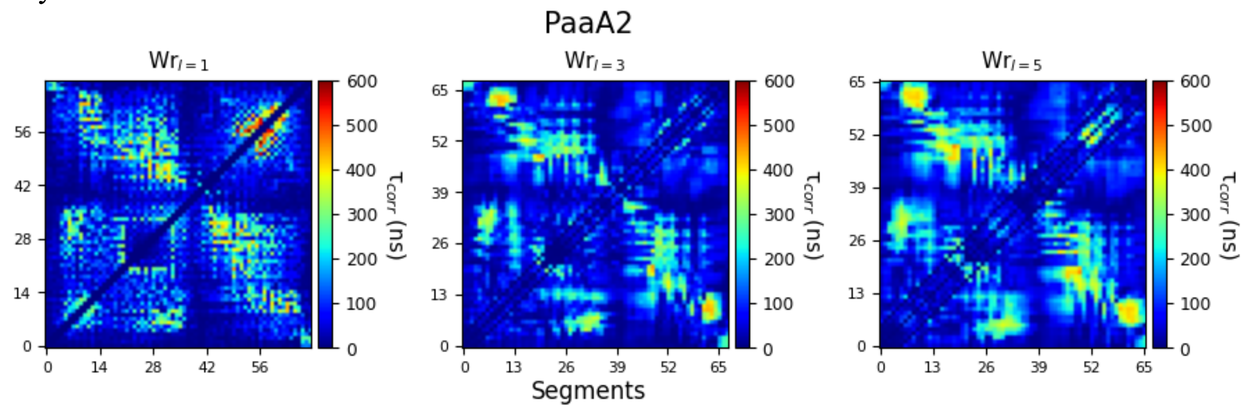


**Supplementary Figure S3-C.** Autocorrelation time ($\tau_{corr}$) matrices for pairwise writhe values between segments, computed at segment lengths of 1, 3 and 5 for a continuous long-timescale molecular dynamics simulation of PaaA2. The autocorrelation times presented here are estimated from integrating the individual autocorrelation functions of each writhe feature up to the time that they first cross zero.

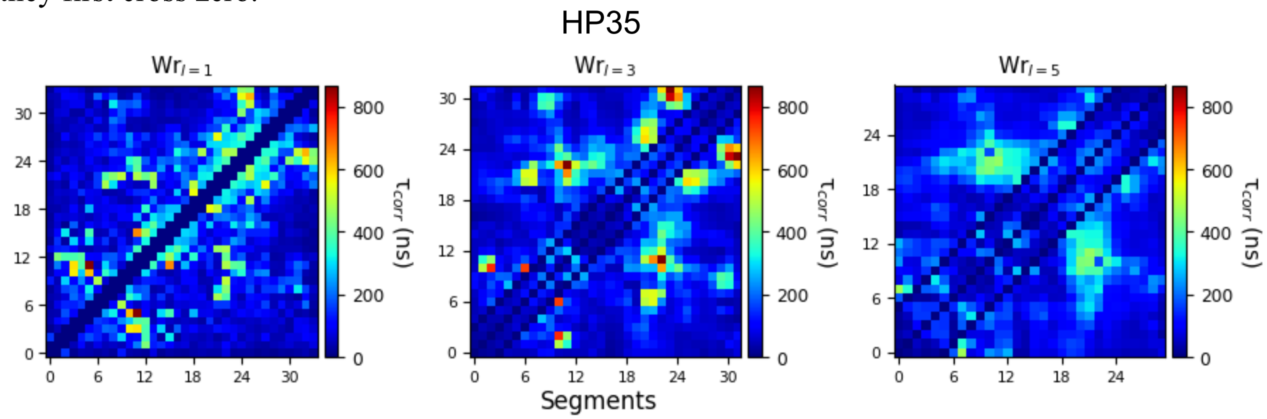


**Supplementary Figure S3-D.** Autocorrelation time ($\tau_{corr}$) matrices for pairwise writhe values between segments, computed at segment lengths of 1, 3 and 5 for a continuous long-timescale molecular dynamics simulation of HP35. The autocorrelation times presented here are estimated from integrating the individual autocorrelation functions of each writhe feature up to the time that they first cross zero.

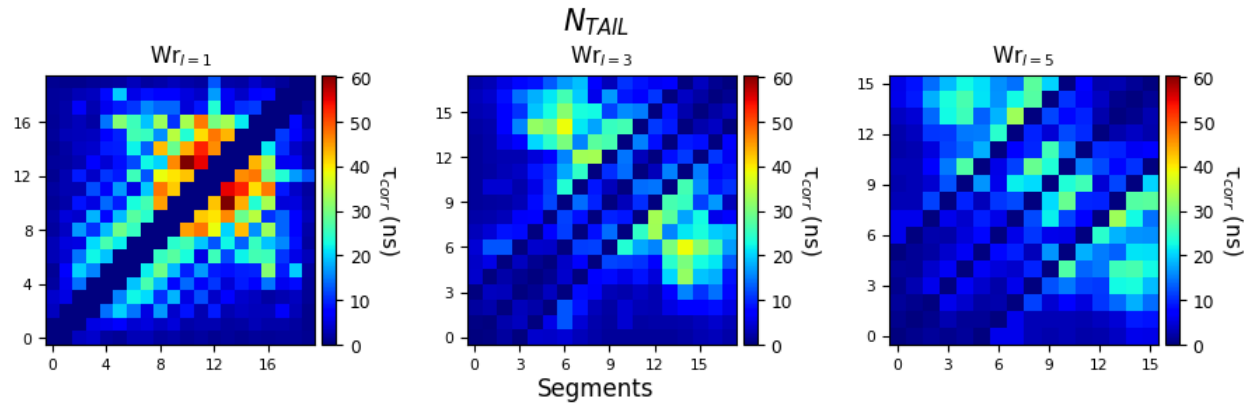


**Supplementary Figure S3-E.** Autocorrelation time ($\tau_{corr}$) matrices for pairwise writhe values between segments, computed at segment lengths of 1, 3 and 5 for a continuous long-timescale molecular dynamics simulation of N$_{TAIL}$. The autocorrelation times presented here are estimated from integrating the individual autocorrelation functions of each writhe feature up to the time that they first cross zero.

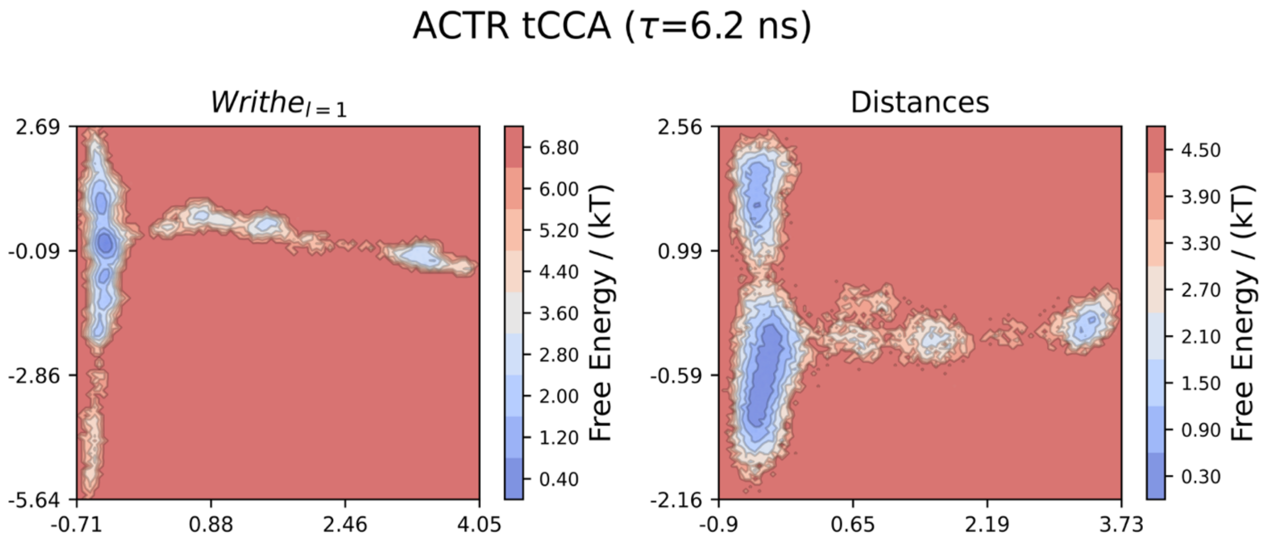


**Supplementary Figure S4. Time-lagged canonical correlation analysis of a long-timescale molecular dynamics simulation of the intrinsically disorder protein, ACTR .** We compare free energy surfaces obtained from performing time-lagged canonical correlation analysis (tCCA) on writhe features computed at segment length 1 (**left**) and inter-residue Euclidean distances (**right**) from a continuous, unbiased 30 μs molecular dynamics simulation of ACTR.

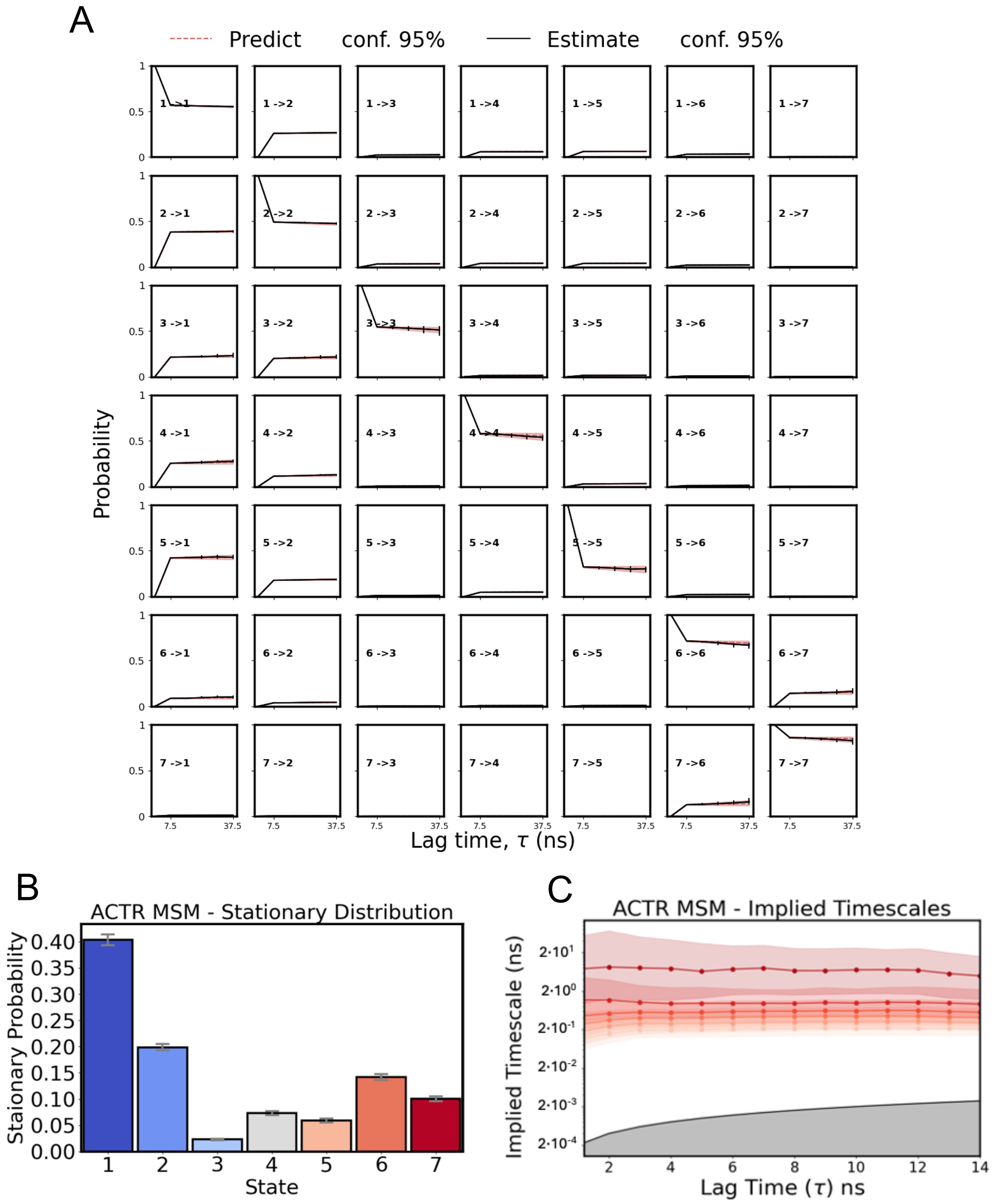


**Supplementary Figure S5**. **ACTR MSM validation tests and stationary distribution. (A)** The Chapman-Kolmogrov (CK) test calculated for the MSM of ACTR constructed from multiscale writhe features computed using segment lengths 1, 3 and 5 ($Wr_{l=1,3,5}$). Writhe features were projected onto a 3-dimensional tCCA space and clustered into 40 microstates using the Kmeans algorithm. The CK test evaluates the dependence of an MSMs predictions on the chosen lag time by comparing the evolution of transition probabilities for each state, i, to every other state, j, for integer multiples of the initial lag time of the model (6 ns). Here, we perform the CK test in the space of 7 macrostates obtained from PCCA++ spectral clustering. Red dotted lines ("Predict")

represent transition probabilities predicted by propagating the transition matrix and the solid black lines (“Estimate”) indicate transition probabilities obtained from transition matrices resampled from the trajectory data at integer multiples of the lag time. The red shaded region indicates the 95% confidence interval of the mean obtained from Gibbs sampling with 1000 samples. **(B)** The stationary distribution for each state of the MSM with error bars showing the deviation of the 95% confidence interval of the mean. **(C)** The log scaled implied timescales obtained from the MSM transition matrices estimated at increasing lag times. The colored shaded regions show the deviation of the 95% confidence interval of the mean of the implied timescales estimated for each lag time. The solid black line and gray shaded region indicate time scales equal to or less than the lag time and represents the threshold for the fastest ITS that can be resolved by the model.

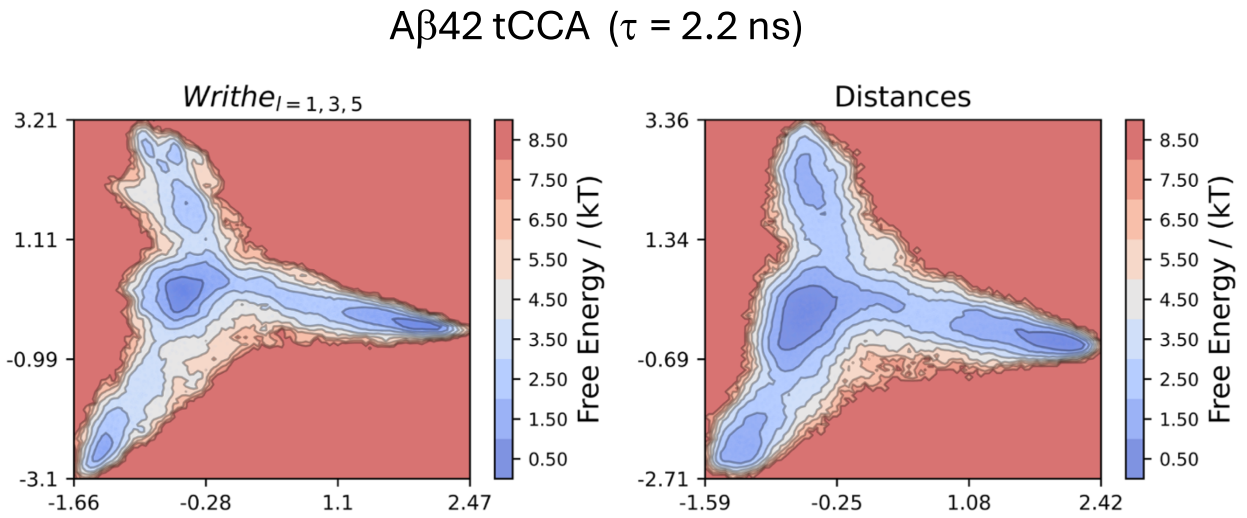


**Supplementary Figure S6. Time-lagged canonical correlation analysis of a long-timescale molecular dynamics simulation of the intrinsically disorder protein, Aβ42.** We compare 2D free energy surfaces obtained from performing time-lagged canonical correlation analysis (tCCA) on multiscale writhe features computed using segment lengths 1, 3 and 5 (**right**) and inter-residue distances (**left**) from 315 μs of molecular dynamics simulation of Aβ42.

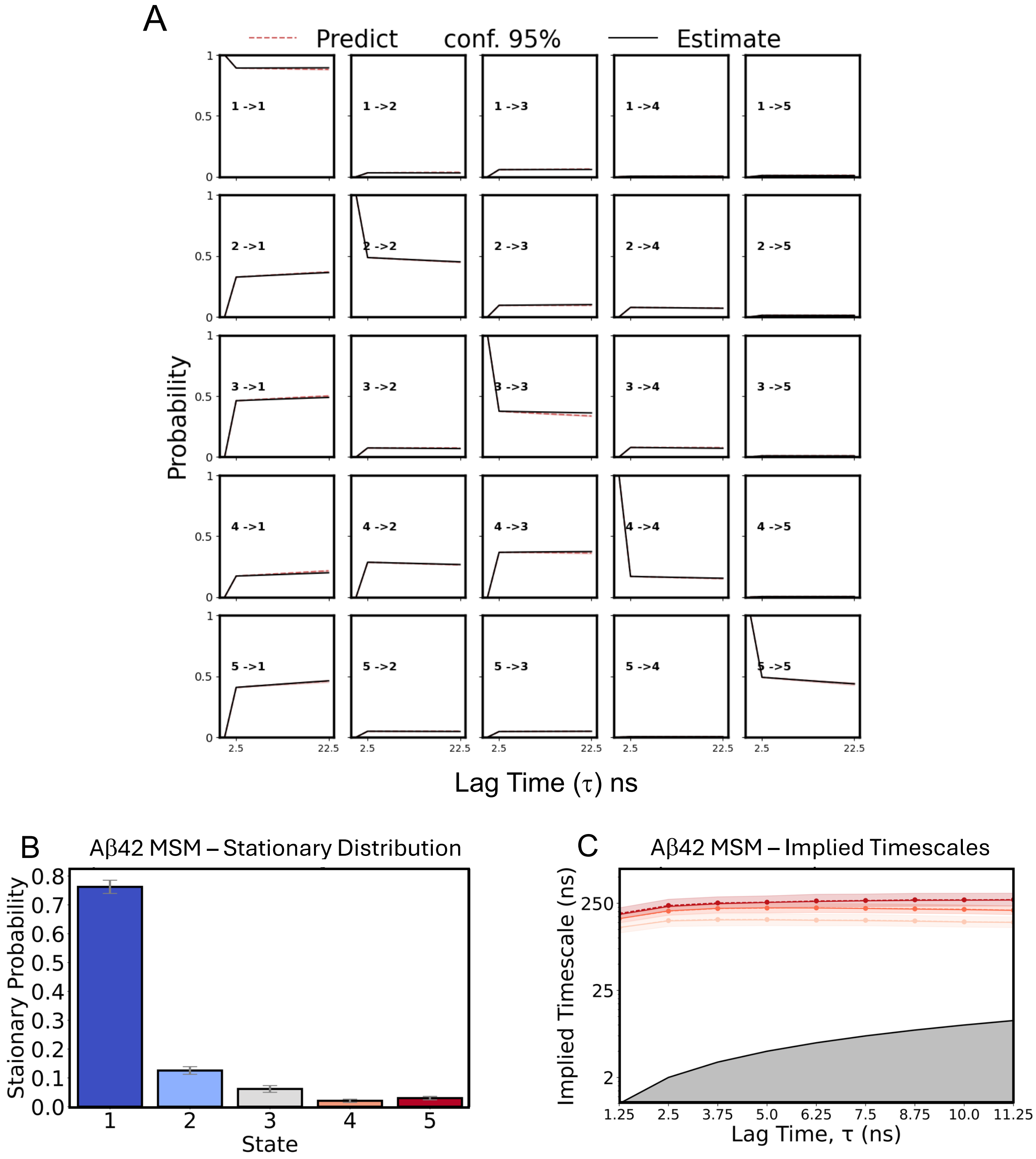


**Supplementary Figure S7**. **Aβ42 MSM validation tests and stationary distribution built from multiscale writhe features. (A)** The Chapman-Kolmogrov (CK) test for the MSM of Aβ-42 constructed from multiscale writhe features computed using segment lengths 1, 3 and 5 ($Wr_{1,3,5}$). Writhe features were projected into a 3-dimensional tCCA space and clustered into 40 microstates using the Kmeans algorithm. The CK test evaluates the dependence of an MSMs predictions on the chosen lag time by comparing the evolution of transition probabilities for each state, i, to every other state, j, for integer multiples of the initial lag time of the model (2.5 ns). Here, we perform the CK test in the space of 5 macro-states obtained from PCCA++ spectral clustering. Red dotted lines ("Predict") represent transition probabilities predicted by propagating the transition matrix and the solid black lines ("Estimate") indicate transition probabilities obtained from transition matrices resampled from the trajectory data at integer multiples of the lag time. The red shaded region indicates the 95% confidence interval of the mean obtained from Gibbs sampling with 1000 samples. **(B)** The stationary distribution for each state of the MSM with error bars showing the deviation of the 95% confidence interval of the mean. **(C)** The log scaled implied timescales obtained from the MSM transition matrices estimated at increasing lag times. The colored shaded regions show the deviation of the 95% confidence interval of the mean of the implied timescales estimated for each lag time. The solid black line and gray shaded region indicate time scales equal to or less than the lag time and represents the threshold for the fastest ITS that can be resolved by the model.

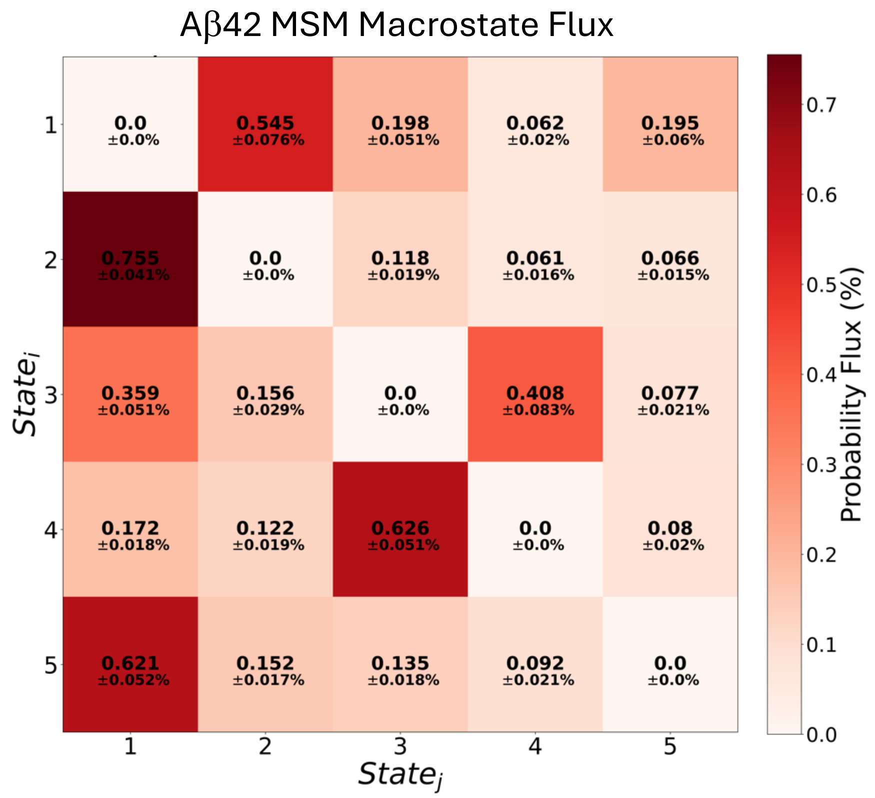


**Supplementary Figure S8. Aβ42 MSM macrostate probability flux matrix for lag time τ = 2.5 ns**. The matrix of pairwise total probability fluxes between metastable sets determined from PCCA++ spectral clustering of a 40 microstate MSM of Aβ42 constructed from multiscale writhe features computed from segment lengths 1, 3 and 5 ($Wr_{1,3,5}$) and estimated at lag time, τ = 2.5 ns. The probability flux describes the flow of probability mass and can be interpreted as the number of transition events along a certain pathway per time unit. Here, fluxes are normalized by the total outgoing flux from each state. The flux matrix shown here is the bootstrap mean of results obtained from Gibbs sampling of the corresponding MSM transition matrix using 1000 samples. Errors report the mean of the upper and lower deviations of the 95% confidence interval of the bootstrap mean.

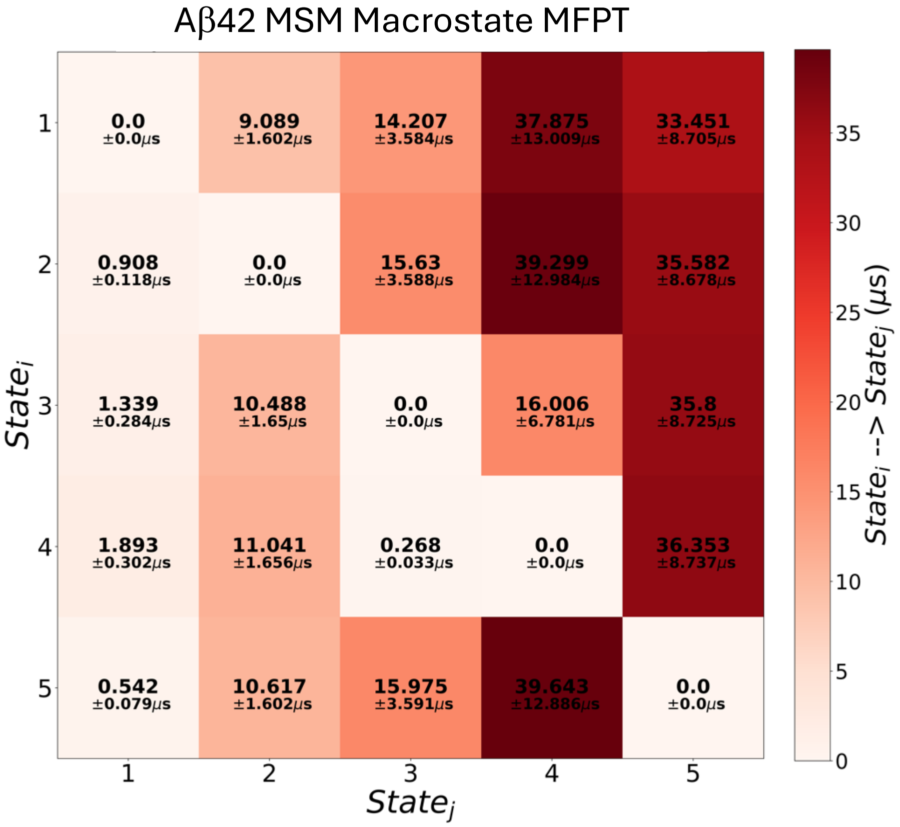


**Supplementary Figure S9. Aβ42 MSM mean first passage time matrix.** The mean first passage time (MFPT) matrix shows the average time duration it takes to transition between metastable sets of microstates in the MSM of Aβ42 constructed from multiscale writhe features computed from segment lengths 1, 3 and 5 ($Wr_{l=1,3,5}$). We compute the mean first passage times using the transition matrix and stationary probabilities of the Aβ42 MSM estimated at lag time, $\tau$ = 2.5 ns. Errors report the mean of the upper and lower deviations of the 95% confidence interval calculated from Gibbs sampling of the transition matrix using 1000 samples.

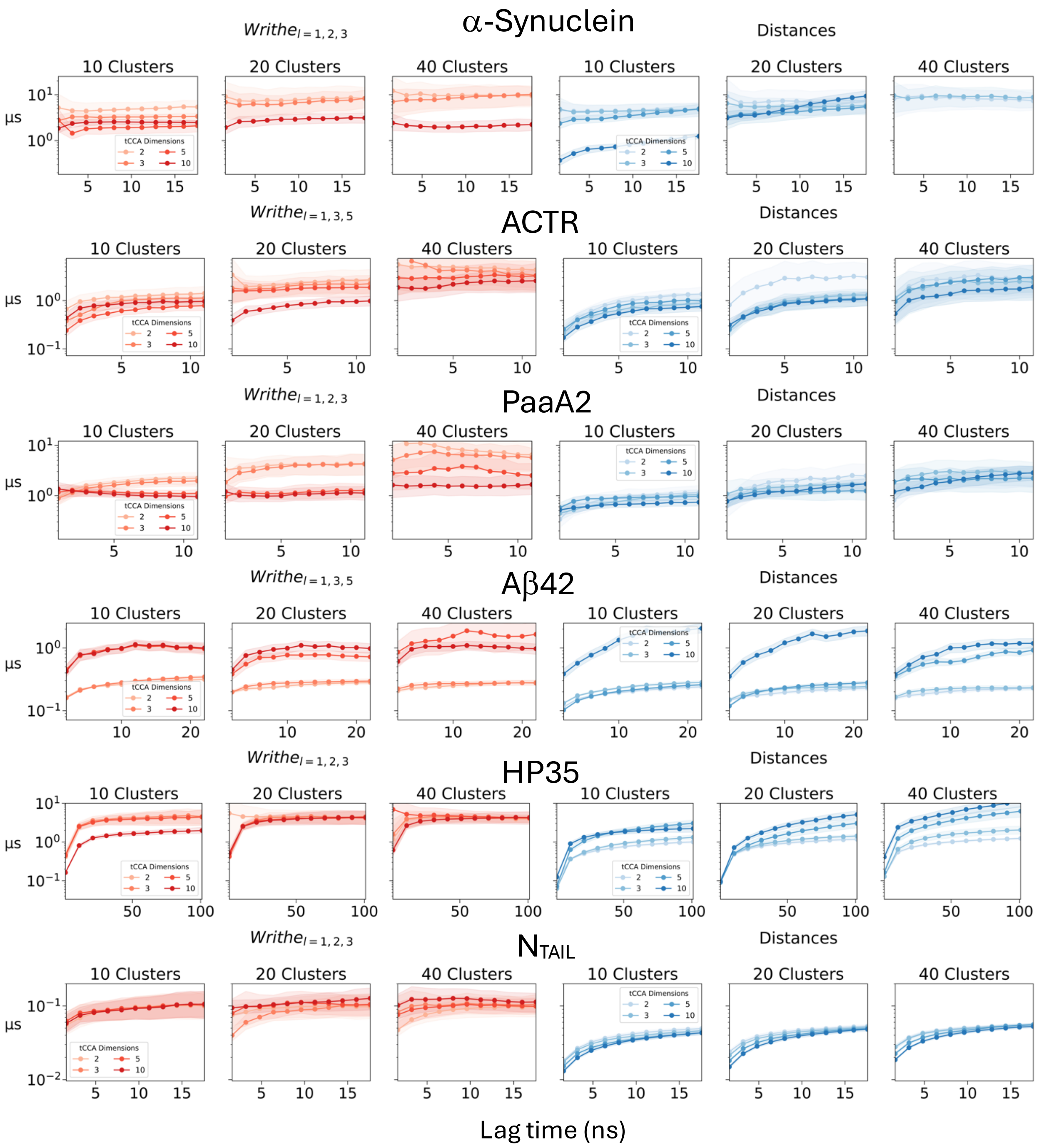
α-Synuclein
ACTR
PaaA2
Aβ42
HP35
NTAIL
Writhe l = 1, 2, 3
Writhe l = 1, 3, 5
Distances
10 Clusters
20 Clusters
40 Clusters
tCCA Dimensions
2
3
5
10
μs
Lag time (ns)

**Supplementary Figure S10. Longest Implied timescales of MSMs as a function of the tCCA dimension and number of clusters for writhe features and inter-residue Euclidean distances.** The longest implied timescale (ITS) of an MSM indicates the relaxation time of the slowest dynamical process it captures. We compare the longest ITS estimated over a range of lag times for MSMs built from the writhe features (red) with the largest kinetic variance (Supplementary Figure S2) in addition to Euclidean distances (blue), for **(A)** α-synuclein, **(B)** ACTR, **(C)** PaaA2, **(D)** Aβ-42, **(E)** HP35 and **(F)** $N_{TAIL}$. After determining the optimal writhe feature set, the MSMs show here are built from clustering tCCA projections using a variable number of K-means clusters and tCCA dimensions. Here, we do not explicitly enforce that MSMs be reversible in order to minimize the impact of estimator error on predicted timescales. Plots excluding specific combinations of tCCA dimensions and clusters failed in estimation due to spare transitions / disconnected state spaces or numerical instability over short changes in the lag time.

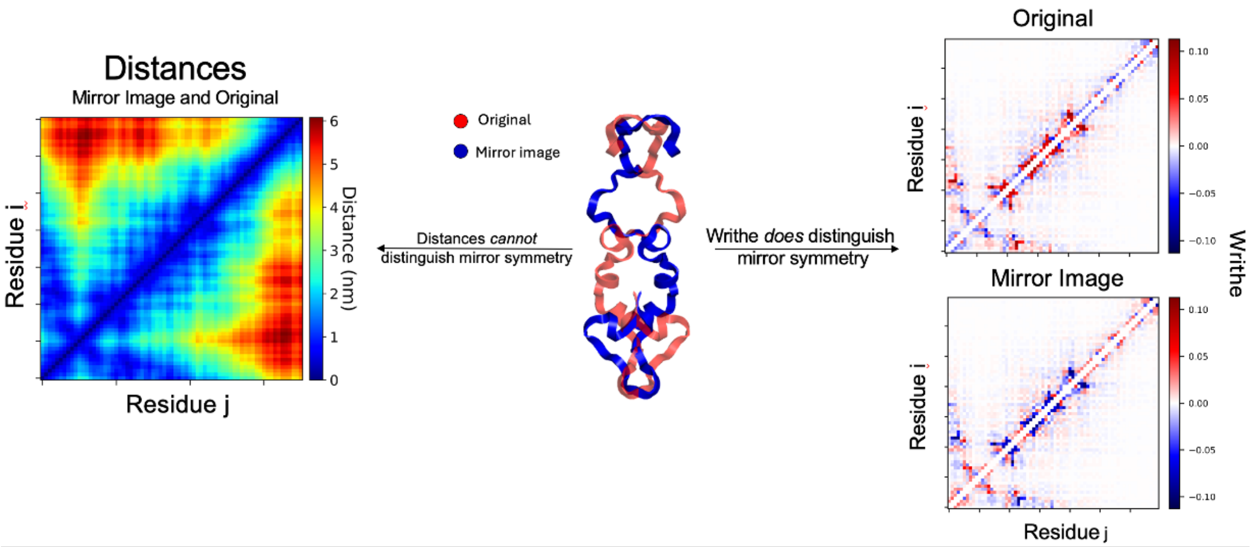


**Supplementary Figure S11. Properties of writhe and Euclidean distance representations.** The same Euclidian distance matrix corresponds to two structures (mirror images) which cannot be made equivalent by rotations and translations. This is because the set of all Euclidian distances for a structure is invariant to actions of the E(3) group. Thus, rotations, translations and parity transformations cannot be recovered from this representation and conformations differing by any of these transformations cannot be distinguished by Euclidian distances. This result can be shown in a rigorous mathematical sense via the relation between inner products and Euclidian distances, and Cholesky decomposition or equivocally by well-established techniques in multidimensional scaling. In contrast, the writhe matrix (matrix of all pairwise segment crossings) differentiates mirror conformations predictably (as seen in the rightmost column) where all features are distinguished by a sign flip. More formally, this is a demonstration of the writhe's equivariance to parity.

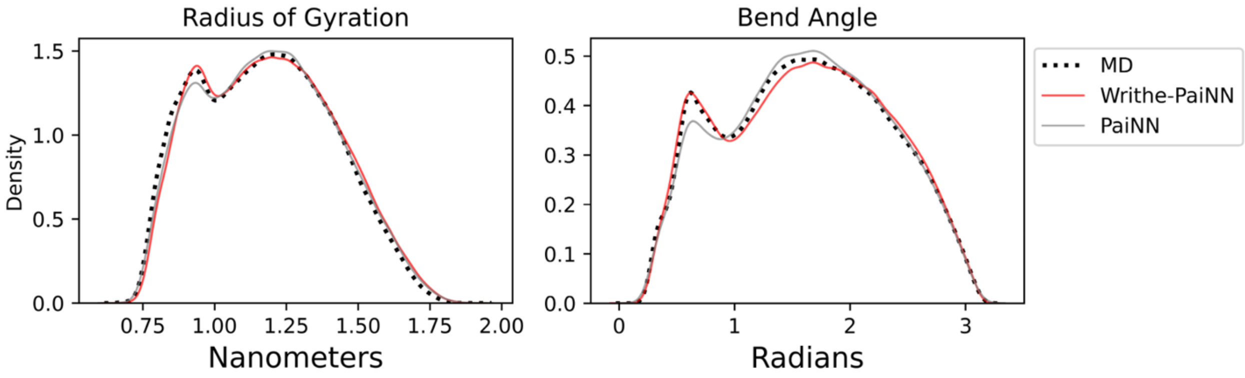


**Supplementary Figure S12. Distributions of parity invariant observables obtained from MD simulation and generative models.** Distributions are obtained from smoothed histograms of the radius of gyration ($R_g$, left) and the angle formed by the first, tenth and last (twentieth) Cα atoms of the 20 residue αsynuclein fragment (right) computed directly from simulation and artificially generated atomic coordinates.

**Appendix A: Numerical computation of the writhe and algorithms**

The numerical computation of the writhe is characterized by the angles, $\theta_{i,j}$, formed at each vertex of the spherical quadrilateral as shown in Figure 2. The sum of these angles minus $\frac{\pi}{2}$ is equal to the surface area of the spherical quadrilateral and is referred to as the *spherical excess*. Here, we take $\vec{a}$, $\vec{b}$, and $\vec{c}$ to be normalized vectors in $\mathbb{R}^3$that represent any set of 3 view direction vectors ($\vec{d}_{i,j}$) from which we compute the contribution to the writhe from 1 of 4 vertices of the spherical quadrilateral shown in Figure 2. This is equivalent to computing the angle between $\vec{a}$ and $\vec{c}$ as seen from the direction of $\vec{b}$, minus $\frac{\pi}{2}$. The angles ($\theta_{i,j}$ in Figure 2) can be computed from cross products as:

$$\sin\left(\theta - \frac{\pi}{2}\right) = \frac{(\vec{a} \times \vec{b}) \cdot (\vec{b} \times \vec{c})}{||\vec{a} \times \vec{b}||\, ||\vec{b} \times \vec{c}||} \tag{A.1}$$

where $sin^{-1}[(\vec{a} \times \vec{b}) \cdot (\vec{b} \times \vec{c})] = \theta - \frac{\pi}{2}$ specifically for vertices constructed following the same convention as $\vec{d}_{i,j}$ triples in Figure 2.This method of computing $\theta_{i,j}$ in Figure 2 is the most precise when the angles between $\vec{a}$, $\vec{b}$, and $\vec{c}$ are very small (< 1 degree) and should be used under single precision arithmetic constraints. Using double precision arithmetic, we can make the following simplifications to construct a faster algorithm to compute $\theta_{i,j}$. Using Lagrange's triple product identity:

$$(\vec{a} \times \vec{b}) \cdot (\vec{b} \times \vec{c}) = (\vec{a} \cdot \vec{b})(\vec{b} \cdot \vec{c}) - (\vec{a} \cdot \vec{c})(\vec{b} \cdot \vec{b}), \tag{A.2}$$

and the Gram determinant:

$$||\vec{a} \times \vec{b}|| = \sqrt{||\vec{a}||^2||\vec{b}||^2 - (\vec{a} \cdot \vec{b})^2}, \tag{A.3}$$

we can rewrite (1) in terms of dot products:

$$\sin\left(\theta - \frac{\pi}{2}\right) = \frac{(\vec{a} \times \vec{b}) \cdot (\vec{b} \times \vec{c})}{||\vec{a} \times \vec{b}||\, ||\vec{b} \times \vec{c}||} = \frac{(\vec{a} \cdot \vec{b})(\vec{b} \cdot \vec{c}) - (\vec{a} \cdot \vec{c})}{\sqrt{\left(1 - (\vec{a} \cdot \vec{b})^2\right)\left(1 - (\vec{b} \cdot \vec{c})^2\right)}}, \tag{A.4}$$

where we used $|| \cdot || = 1$ , $\vec{b} \cdot \vec{b} = ||\vec{b}||^2$ .We can compute the surface area (*SA*) shown in Figure 2 by evaluating (1) or (4) on the view direction vectors ($\vec{d}_{i,j}$) defining each vertex, taking the $sin^{-1}$and summing over all angles, $\theta_{i,j}$. Using equation (2) allows us to writhe the surface area (*SA*) in terms of only 6 unique dot products which we denote with Greek letters in the following:

$$\begin{aligned} &\alpha = \vec{d}_{1,3} \cdot \vec{d}_{2,3}, \quad \beta = \vec{d}_{1,3} \cdot \vec{d}_{1,4} \\ &\gamma = \vec{d}_{1,4} \cdot \vec{d}_{2,3}, \quad \sigma = \vec{d}_{1,4} \cdot \vec{d}_{2,4} \\ &\mu = \vec{d}_{1,3} \cdot \vec{d}_{2,4}, \quad \phi = \vec{d}_{2,3} \cdot \vec{d}_{2,4} \end{aligned} \implies \begin{aligned} SA =& sin^{-1}\frac{\alpha \cdot \beta - \gamma}{\sqrt{(1-\alpha^2)(1-\beta^2)}} + sin^{-1}\frac{\beta \cdot \sigma - \mu}{\sqrt{(1-\beta^2)(1-\sigma^2)}} \\ &+ sin^{-1}\frac{\phi \cdot \alpha - \mu}{\sqrt{(1-\phi^2)(1-\alpha^2)}} + sin^{-1}\frac{\sigma \cdot \phi - \gamma}{\sqrt{(1-\sigma^2)(1-\phi^2)}} \end{aligned} \tag{A.5}$$

Regardless of which approach is used to compute the surface area element (*SA*), the derivation of the writhe for a single crossing is completed with the same factor, $\delta_{\pm}$, which accounts for the sign of the crossing and takes on a value of 1 or -1. The sign factor can be computed from the determinant or scalar triple product of the two crossing segments, $\vec{s}_1$, $\vec{s}_2$, and the view direction vector, $\vec{d}_{1,3}$: $\delta_{\pm} = \text{sign}(\vec{d}_{1,3} \cdot (\vec{s}_2 \times \vec{s}_1)) = \text{sign}(det|\vec{d}_{1,3}\ \vec{s}_2\ \vec{s}_1|)$. From these definitions, the writhe of a crossing is:

$$\mathrm{W_r} = \frac{\delta_{\pm}}{2\pi} \cdot SA \tag{A.6}$$

The normalization constant for each individual writhe value, $2\pi$, can be understood as $2/4\pi$ where $4\pi$ normalizes the area of the unit sphere and the factor of 2 considers that a crossing seen from one direction is also seen from the antipodal direction. Both approaches presented here are analytically correct and equivalent, however, minor discrepancies arise from floating point errors in numerical computation. Consequently, both algorithms are available in the python package accompanying this work.

| Method | Dataset | Mean wall-clock time $\pm\,\sigma$ (ms) | Number of trials |
|---|---|---|---|
| Dot products | 1 structure<br>140 residues<br>9,453 crossings | $3.14 \pm 0.066$ | 10,000 |
| Cross products | | $6.82 \pm 0.093$ | |

**Supplementary Table 1**. **Wall-clock times to compute the writhe between all segment pairs of a single structure**. The writhe between all segment pairs was computed for a single conformation of full-length a-synuclein using a segment length of 1. Both algorithms are implemented in python using *PyTorch* and utilize broadcasting to compute the writhe of all segment pairs simultaneously. Given the number of residues in a structure, $n$, and the segment length, $l$, the number of non-trivial crossings between segments can be computed as $\frac{(n-l)^2-(n-l)-2(n-2l)}{2}$. Here, "non-trivial" refers to segment pairs that do not systematically give a writhe of zero.

### Appendix B: Definition of atom-wise writhe features

To incorporate writhe into atom-wise graph neural network message-passing operations, we compute cumulative writhe features between atoms using the writhe graph Laplacian defined below. The writhe graph Laplacian operates on the graph implied by the segments (edges) connecting pairs of atoms (nodes) used to compute the writhe. The definition of the discrete computation of the writhe used in this work always implicitly defines a corresponding graph Laplacian. This allows us to systematically obtain pair-wise writhe features between atoms from pair-wise computation of the writhe between segments connecting atoms.

Here, we consider only the writhe computed at segment length 1. Thus, for n atoms, there are ($n$ - 1) segments. The atom-segment relationships can be encoded using the incidence matrix, $B \in \mathbb{R}^{n\times(n-1)}$, where rows represent atoms and columns represent segments. This incidence matrix corresponds to a graph comprised of nodes sequentially connected by undirected edges. In general, the incidence matrix, $B$, takes the form:

$$B = \begin{bmatrix} 1 & 0 & 0 & \cdots & 0 \\ 1 & 1 & 0 & \cdots & 0 \\ 0 & 1 & 1 & \cdots & 0 \\ 0 & 0 & 1 & \cdots & 0 \\ \vdots & \vdots & \vdots & \ddots & 0 \\ 0 & 0 & 0 & 0 & 1 \end{bmatrix} \tag{B.1}$$

Pairwise computation of the writhe between segments yields the symmetric writhe matrix, $W \in \mathbb{R}^{(n-1)\times(n-1)}$:

$$W = \begin{bmatrix} 0 & 0 & w_{1,3} & w_{1,4} & \cdots & w_{1,n-1} \\ 0 & 0 & 0 & w_{2,4} & \cdots & w_{2,n-1} \\ w_{1,3} & 0 & 0 & 0 & \cdots & w_{2,n-1} \\ w_{1,4} & w_{2,4} & 0 & 0 & \cdots & w_{4,n-1} \\ \vdots & \vdots & \vdots & \vdots & \ddots & \vdots \\ w_{1,n-1} & w_{2,n-1} & w_{3,n-1} & w_{4,n-1} & \cdots & 0 \end{bmatrix} \tag{B.2}$$

where each $w_{i,j}$ represents the scalar value of the writhe between segments ($i, j$) and

$$w_{ij} = \begin{cases} 0, & \text{if } |i - j| < 2 \\ w_{ji}, & \text{for all } i, j \end{cases} . \tag{B.3}$$

The graph Laplacian[7], $L_{wr} \in \mathbb{R}^{n\times n}$, corresponding to the incidence matrix, B, weighted by the writhe matrix, $W$, is defined as:

$$L_{wr} = BWB^T \tag{B.4}$$

and takes the general form:

$$L_{wr} = \begin{bmatrix} 0 & 0 & w_{1,3} & (w_{1,3} + w_{1,4}) & \cdots & w_{1,n-1} \\ 0 & 0 & w_{1,3} & (w_{1,3} + w_{1,4} + w_{2,4}) & \cdots & (w_{1,n-1} + w_{2,n-1}) \\ w_{1,3} & w_{1,3} & 0 & w_{2,4} & \cdots & (w_{2,n-1} + w_{3,n-1}) \\ (w_{1,3} + w_{1,4}) & (w_{1,3} + w_{1,4} + w_{2,4}) & w_{2,4} & 0 & \cdots & (w_{3,n-1} + w_{4,n-1}) \\ \vdots & \vdots & \vdots & \vdots & \ddots & \vdots \\ w_{n-1,1} & (w_{1,n-1} + w_{2,n-1}) & (w_{2,n-1} + w_{3,n-1}) & (w_{3,n-1} + w_{4,n-1}) & \cdots & 0 \end{bmatrix}$$

Each index in $L_{wr}$ is the sum over all segment crossings in which a pair of atoms both appear and are not a part of the same segment. Programmatically, the operations described above are performed using summation operations on graphs rather than directly evaluating matrix operations. This allows us to avoid redundant computations involving symmetric matrices and generalize the writhe graph Laplacian to the case where each $w_{ij}$ is a vector ($\vec{w}_{i,j}$).

To demonstrate the similarity of a symmetric writhe matrix and the writhe graph Laplacian we compare the average $Wr_{l=1}$ writhe matrix computed from a 30μs MD simulation of ACTR with the average writhe graph Laplacian for the same trajectory in Supplementary Figure S14.

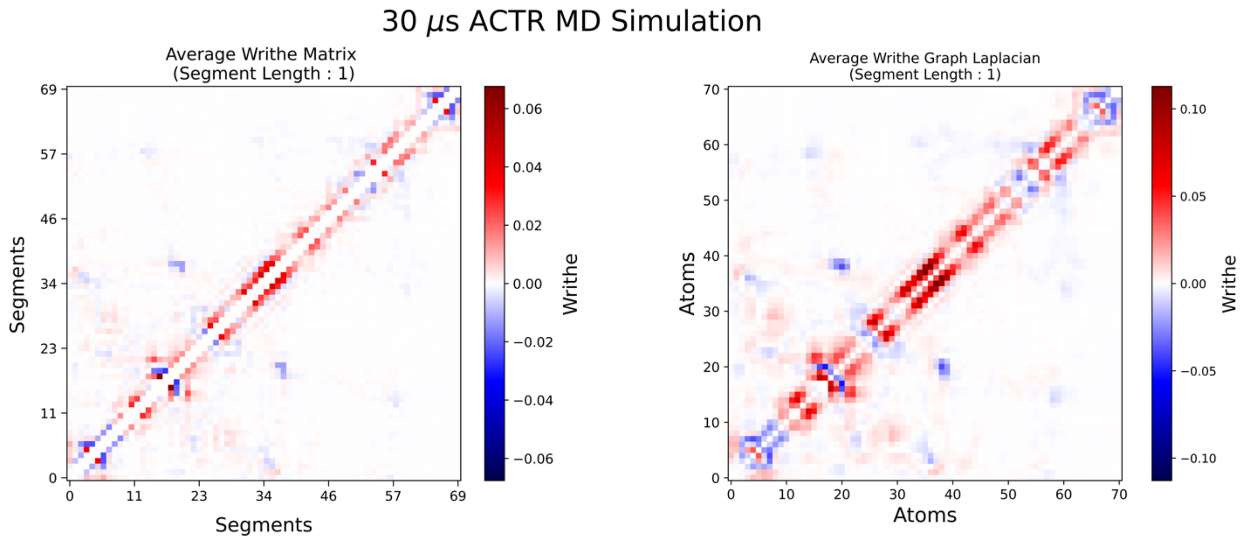


**Supplementary Figure S13: Comparison of the average $Wr_{l=1}$ writhe matrix and the writhe-graph Laplacian computed from a 30 μs MD simulation of ACTR.**

### Appendix C: PaiNN architecture implementation and DDPM training

All DDPMs were trained on structural data obtained from a 100 μs all atom molecular dynamics simulation of a 20 residue C-terminal fragment of α-synuclein[8] using the a99SB-disp protein force field[9] and a99SB-disp water model.[9] We train DDPMs on the full simulation dataset comprised of ≈550,000 structures and consider only the coordinates of Cα atoms to minimize computational expense. Both the PaiNN and Writhe-PaiNN architectures have input blocks that assign atoms and bonds trainable embedding vectors (categorical encodings) that are treated as invariant scalar features ($s_i$) and compute direction vectors ($\vec{r}_{i,j}$) and Euclidean distances ($\|\vec{r}_{i,j}\|$) between atoms $(i, j)$. In the writhe model, the input block also computes pair-wise vector and scalar writhe features (Appendices A and B). The dimension of the model is a hyperparameter that determines the dimension of categorical embeddings (atom-types and bonds) and positional encodings (distances and writhe when applicable). Equivariant vector features ($v_i$) are lifted to the dimension of the model by repeating each vector (a set of vectors for each atom or node) and multiplying each copy by a unique, learned scalar. Here, we use a model dimension of 64 in all experiments. We repeat message passing layers 8 times to construct the message passing blocks for both the PaiNN and Writhe-PaiNN. Both models are equipped with the same output block that transforms higher dimension invariant scalar ($s_i$) and equivariant vector ($v_i$) features into a prediction of the *score field*, which has the same dimension as the coordinates of the target structure.

We trained models with the ADAM[10] optimizer using a learning rate of 0.0001 until the loss[11] stabilized, after which point we monitored how well generated samples from models were able to reproduce distributions from the training data. For the ≈550,000 frame α-synuclein dataset, which samples a diverse conformational space, we found that both the PaiNN and Writhe-PaiNN models continued to improve agreement with training data distributions until ~750 epochs and did not begin to significantly overfit until ~1200 epochs. For comparison purposes, we present the results from PaiNN and Writhe-PaiNN DDPMs trained for 900 epochs. We reiterate that our goal is to test the symmetry properties and overall efficacy of the proposed writhe-based neural network architecture and compare to the original PaiNN architecture, rather than obtaining a model the generalizes across chemical space and can predict ensembles for arbitrary sequences.

### References


(1) Konstantin, K.; Langowski, J. Computation of writhe in modeling of supercoiled DNA. *Biopolymers* **2000**, *54* (5), 307-317.

(2) Călugăreanu, G. L'intégrale de Gauss et l'analyse des nœuds tridimensionnels. *Rev. Math. Pures Appl.* **1959**, *4*, 5–20.

(3) Røgen, P.; Bohr, H. A new family of global protein shape descriptors. *Mathematical Biosciences* **2003**, *182* (2), 167-181.

(4) Røgen, P.; Fain, B. Automatic classification of protein structure by using Gauss integrals. *Proceedings of the National Academy of Sciences* **2002**, *100* (1), 119-124.

(5) Hotelling, H. Relations Between Two Sets of Variates. *Biometrika* **1936**, *28* (3/4), 321.

(6) Wu, H.; Noé, F. Variational Approach for Learning Markov Processes from Time Series Data. *Journal of Nonlinear Science* **2019**, *30* (1), 23-66.

(7) Strang, G. *Linear algebra and learning from data*; Wellesley-Cambridge Press, 2019.

(8) Robustelli, P.; Ibanez-de-Opakua, A.; Campbell-Bezat, C.; Giordanetto, F.; Becker, S.; Zweckstetter, M.; Pan, A. C.; Shaw, D. E. Molecular Basis of Small-Molecule Binding to α-Synuclein. *Journal of the American Chemical Society* **2022**, *144* (6), 2501-2510.

(9) Robustelli, P.; Piana, S.; Shaw, D. E. Developing a molecular dynamics force field for both folded and disordered protein states. *Proceedings of the National Academy of Sciences* **2018**, *115* (21), E4758-E4766.

(10) Kingma, D. P.; Ba, J. Adam: A Method for Stochastic Optimization. arXiv preprint arXiv:1412.6980: 2014.

(11) Song, Y.; Durkan, C.; Murray, I.; Ermon, S. Maximum Likelihood Training of Score-Based Diffusion Models. *arXiv preprint arXiv:2101.09258* **2021**.
